# Logan: Planetary-Scale Genome Assembly Surveys Life’s Diversity

**DOI:** 10.1101/2024.07.30.605881

**Authors:** Rayan Chikhi, Téo Lemane, Raphaël Loll-Krippleber, Mercè Montoliu-Nerin, Brice Raffestin, Antonio Pedro Camargo, Carson J. Miller, Mateus Bernabe Fiamenghi, Daniel Paiva Agustinho, Sina Majidian, Greg Autric, Maxime Hugues, Junkyoung Lee, Roland Faure, Kristen D. Curry, Jorge A. Moura de Sousa, Eduardo P. C. Rocha, David Koslicki, Paul Medvedev, Purav Gupta, Jessica Shen, Alejandro Morales-Tapia, Kate Sihuta, Peter J. Roy, Grant W. Brown, Robert C. Edgar, Anton Korobeynikov, Martin Steinegger, Caleb A. Lareau, Pierre Peterlongo, Artem Babaian

## Abstract

The breadth of life’s diversity is unfathomable, but public nucleic acid sequencing data offers a window into the dispersion and evolution of genetic diversity across Earth. However the rapid growth and accumulation of sequence data have outpaced efficient analysis capabilities. The largest collection of freely available sequencing data is the Sequence Read Archive (SRA), comprising 27.3 million datasets or 5 × 10^16^ basepairs. To realize the potential of the SRA, we constructed Logan, a massive sequence assembly transforming short reads into long contigs and compressing the data over 100-fold, enabling highly efficient petabase-scale analysis. We created Logan-Search, a *k*-mer index of Logan for free planetary-scale sequence search, returning matches in minutes. We used Logan contigs to identify *>*200 million plastic-degrading enzyme homologs, and validate novel enzymes with catalytic activities exceeding current reference standards. Further, we vastly expand the known diversity of proteins (30-fold over UniRef50), plasmids (22-fold over PLSDB), P4 satellites (4.5-fold), and the recently described Obelisk RNA elements (3.7-fold). Logan also enables ecological and biomedical data mining, such as global tracking of antimicrobial resistance genes and the characterization of viral reactivation across millions of human BioSamples. By transforming the SRA, Logan democratizes access to the world’s public genetic data and opens frontiers in biotechnology, molecular ecology, and global health.

## 1 Main

DNA sequencing has revolutionized our perspective on life’s diversity, yet the majority of the world’s sequencing data are inaccessible to systematic search and analysis. The Sequence Read Archive (SRA) houses over 50 petabases (Pbp; 5.0×10^16^) of public sequencing data, and is growing exponentially (Fig. 1a) [1]. This data represents billions of dollars of global research output, spanning all known life and covering every continent (Fig. 1).

**Figure 1:**
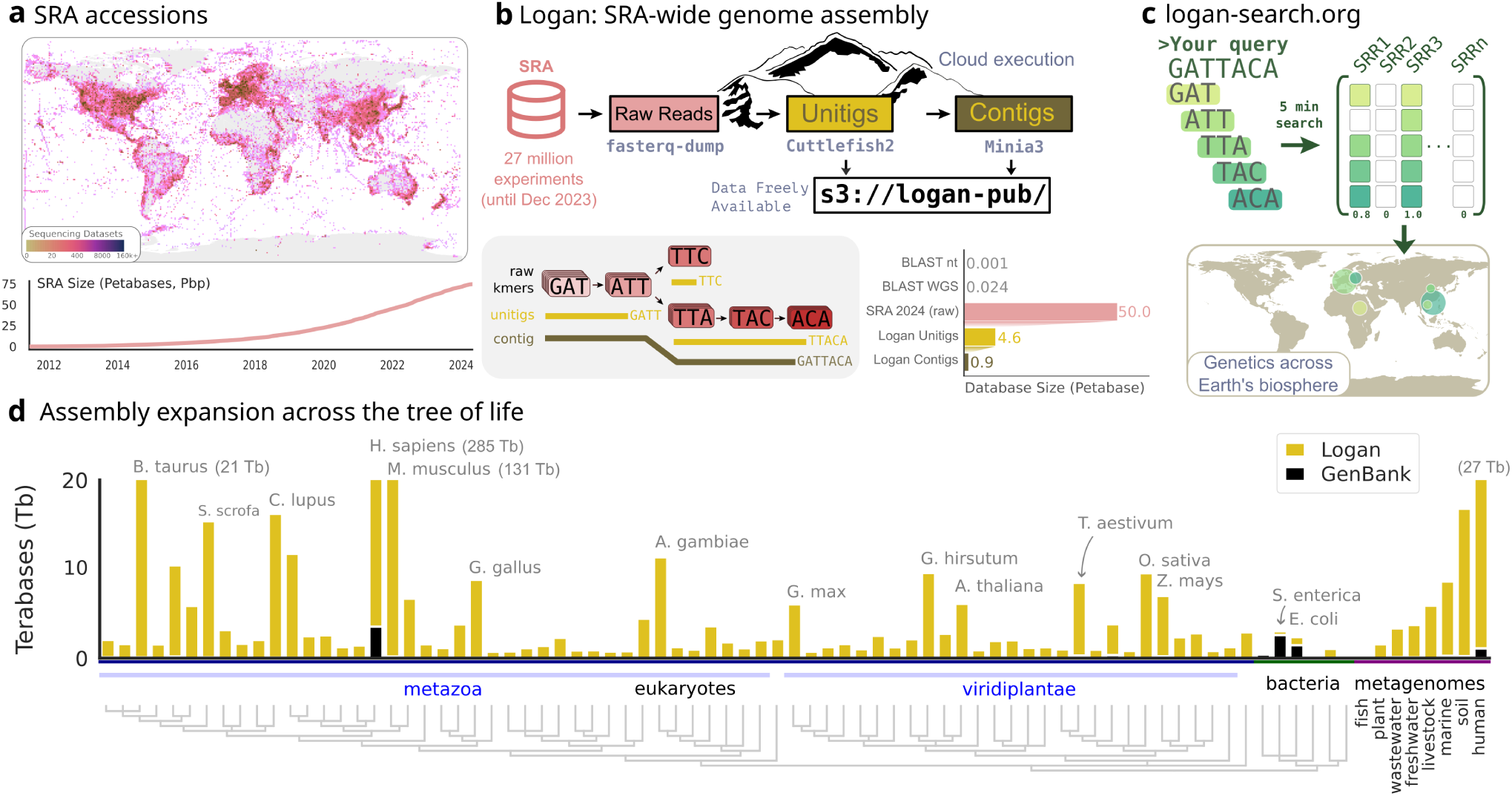
Assembling all accessions of the SRA using a cloud architecture into unitigs and contigs. **(a)** Geographic distribution of samples over the Sequence Read Archive (SRA), and the near-exponential growth of SRA in terms of number of cumulative accession size of raw data. **(b)** Top diagram describes the cloud computation workflow of Logan, starting from SRA reads, then computing unitigs and contigs assemblies, and finally uploading data to our public repository. Bottom left diagram shows a toy dataset with *k*-mers extracted from raw reads, then unitigs and contigs constructed. Bottom right bar plot represents the size of the SRA compared to Logan assembled unitigs and contigs in sum of bases, and WGS and BLAST databases. **(c)** The logan-search.org service enables searching an arbitrary query (example: “GATTACA”) against the full unitig index of the SRA in less than 5 min; hits are mapped to their geographic origins. **(d)** Tree of Life sampled with the 116 most abundant taxa from NCBI GenBank WGS as well as 116 most abundant taxa in Logan assemblies, according to NCBI taxonomy. Black bars represent the total number of assembled bases in GenBank WGS, and yellow bars the additional number of bases in Logan contigs. Bars exceeding 20 terabases are capped and their true total assembly size is annotated. Assembled bases for a subset of metagenome types are represented separately as the 8 rightmost bars.

Analyses of the SRA have yielded profound scientific discoveries, from hundreds of thousands of novel viruses to shifts in antibiotic resistance patterns [2, 3, 4, 5, 6, 7, 8, 9]. Yet the methods for massive-scale sequence analyses, based on assembly or *k*-mer indexing, face computational and economic constraints. The largest collection of public assembled data, NCBI GenBank WGS, spans less than 3% of the SRA [10], while *k*-mer indexing of reads have not scaled beyond 9% of direct SRA data (Table 1).

**Table 1:** Size of existing indexed data vs Logan. The MetaGraph-SRA and Pebblescout-SRA rows refer to all SRA accessions indexed by MetaGraph and Pebblescout respectively [4, 6]. The NCBI SRA row refers to all public accessions from the SRA as of December 10th 2023. The Reads column refers to the number of bases in SRA reads for the considered dataset. The Index/Seqs column indicates the sum of all sub-indices sizes (for Branchwater [8], MetaGraph and Pebblescout) or the size of compressed sequences for all accessions (for SRA and Logan). The “Compress.” column gives the compression ratio between the size of reads (Reads column) as if each base was stored using 8 bits, and the Index/Seqs column.

| Dataset | Type | Reads | # accessions | Index/Seqs | Compress. |
| --- | --- | --- | --- | --- | --- |
| Sourmash Branchwater | Index | 1.4 Pbp | 682,688 | 7 TB | 191× |
| MetaGraph-SRA | Index | 3.3 Pbp | 1,891,328 | 7 TB | 473× |
| Pebblescout-SRA | Index | 3.7 Pbp | 4,141,058 | 171 TB | 21× |
| NCBI SRA (Dec 2023) | Raw data | 50.3 Pbp | 27,764,168 | 19 PB | 3× |
| Logan, Unitigs (v1) | Assembly | 48.3 Pbp | 27,311,279 | 2 PB | 24× |
| Logan, Contigs (v1.1) | Assembly | 44.1 Pbp | 26,788,829 | 315 TB | 140× |
| Logan-Search | Index | 43.9 Pbp | 23,404,655 | 1 PB | 44× |

To enable SRA-wide analysis, we developed Logan, an assembly of 96% of the SRA (27 million accessions as of December 2023) and a suite of associated tools. Using massively parallel cloud processing, we transformed 44.1 petabases of raw sequencing data into 0.9 petabases of long assembled contigs. Logan achieved a *>*100-fold compression of the original SRA data, and is 36-fold larger than GenBank WGS. Logan assembly fundamentally changes the economics and speed of SRA-wide searches. To demonstrate this, we developed Logan-Search, a *k*-mer index for finding a nucleotide sequence across all Logan assemblies in minutes. For protein homology search, we aligned using DIAMOND2 all Logan contigs to a protein query in 11 wall-clock hours, with a near 20-fold cost decrease relative to previous methods such as Serratus [11, 2].

We demonstrate Logan’s utility by making discoveries in three domains. First, for bioprospecting, we performed an SRA-wide search for homologs of plastic-degrading enzymes with sensitivity to *≈* 40% amino acid sequence identity, discovering over 200M novel enzymes, including several which we validated as having higher and more varied catalytic activities than previous standards. Second, for scalable clinical discovery, we used Logan-Search to screen millions of human datasets for viral gene expression sequences. We uncovered recurrent Human Herpesvirus-6 reactivation in tumor-infiltrating lymphocyte therapy products, broadening the characterization of viral reactivation in cell therapies *ex vivo* [12]. Third, we mined Logan contigs for proteins, plasmids and subviral elements. This effort massively expanded the known protein universe, with a 30-fold increase in protein diversity over UniRef at 50% amino-acid clustering identity. We also obtained a 22-fold increase in plasmid families, a 4.5-fold increase in the diversity of P4 satellites, and a 3.7-fold expansion of the recently described Obelisk-like species [9].

### A Petabyte-Scale Assembly and Search Engine of the Sequence Read Archive

To construct the Logan assemblage we applied two complementary assembly strategies to the entirety of the SRA (as of 2023-12-10), comprising 27.3 million SRA accessions (Fig. 1b). The first strategy generates unitigs, which are near-lossless representations aiming to preserve sequence content from a sample, making them ideal for sensitive *k*-mer-based search. The second strategy builds on the unitigs to create contigs, which form longer, consensus sequences by resolving small biological variations. Contigs are optimized for protein identification and other downstream analyses.

In total, the Logan assemblage is 4.59 Pbp of unitigs (2.18 petabytes compressed) and 0.90 Pbp of contigs (0.31 petabytes compressed, Fig. 1b), generated in approximately 30 hours of wall-clock time, using a peak of 2.18 million CPU cores (Methods, Extended Data Fig 1). All assemblies and documentation are publicly hosted and freely available (s3://logan-pub/, https://github.com/IndexThePlanet/Logan). To make the Logan assemblage rapidly search-able we developed Logan-Search, a 1 petabyte *k*-mer (k=31) index of Logan unitigs from 23.4M SRA accessions. For queries of up to 1 kb, Logan-Search returns all indexed SRA accessions containing a user-set fraction of the query *k*-mers, along with associated SRA metadata. This provides rapid insights into the environmental and geographic distributions of a gene (Fig. 1c). Logan-Search surpasses previous non-Logan based SRA sequence-search efforts [8, 4, 6] by at least an order of magnitude in both the number of accessions and total coverage (Table 1). To democratize SRA-scale sequence exploration, we deployed Logan-Search as a web interface (https://logan-search.org), enabling researchers to freely query Logan unitigs in a few minutes.

Logan’s 0.9 petabases of contigs span the entire tree of life, providing orders of magnitude more assembled data for nearly every sequenced species (Fig. 1d). For key model organisms and agricultural species, the amount of assembled data increases dramatically, rising from less than 4 Tbp in GenBank to approximately 285 Tbp for *Homo sapiens* (70-fold increase), from 0.9 Tbp to 21 Tbp for *Bos taurus* (cattle, 23-fold increase), from 0.5 Tbp to 8.8 Tbp for *Gallus gallus* (chicken, 18-fold increase), and from 0.2 Tbp to 7 Tbp for *Zea mays* (maize, 35-fold increase).

The assembled metagenome samples in Logan exceed 130 terabases (over 5.7 million accessions), 144-fold more bases than a comparable collection of short read assembled metagenomes (MGNify [13]). To quantify the novel sequence content within Logan’s metagenomes, we used sketching to estimate the number of distinct 31-mers and compared this to the entirety of the GenBank WGS database. Logan’s metagenome collection contains an estimated 33.9 trillion distinct *k*-mers that appear two or more times in an accession, a nearly 4-fold increase over the 8.7 trillion found in all of GenBank WGS. Crucially, of Logan’s 33.9 trillion metagenomic *k*-mers, 32.3 trillion (95%) are not present anywhere in the GenBank WGS database, highlighting a vast reservoir of previously uncharacterized genetic diversity.

### Expanding the arsenal of plastic-active enzymes

Logan is a powerful tool for enzyme discovery. Since the mid-20^th^ century humans have manufactured in excess of 12 gigatons of plastics, with the majority ending up as waste which degrades into micro- and nanoplastics. These particles infiltrate global ecosystems and food supplies, where they bio-accumulate to high incidence in humans [14, 15]. The 2016 discovery of a polyethylene terephthalate (PET) plastic degrading enzyme in *Ideonella sakaiensis* (*Is*PETase) has catalyzed bio-prospecting and bioengineering efforts to identify novel and high-efficiency enzymes for recycling and remediating plastic polymers [16, 17, 18, 19, 20].

To expand the diversity of plastic-active enzymes, we created a search query from the 213 validated plastic-active enzyme sequences in the PAZy database [18]. These sequences group into 11 CATH protein domains [21], showing polyphyletic activity against varying plastic substrates (Fig. 2a). First, we searched the 213 sequences in the NCBI nr database (0.003 Pbp, DIAMOND2 blastp) and recovered 2.73 million distinct sequences or 1.05 million non-redundant enzyme homologs (clustered at 90% amino acid identity, aaid).

**Figure 2:**
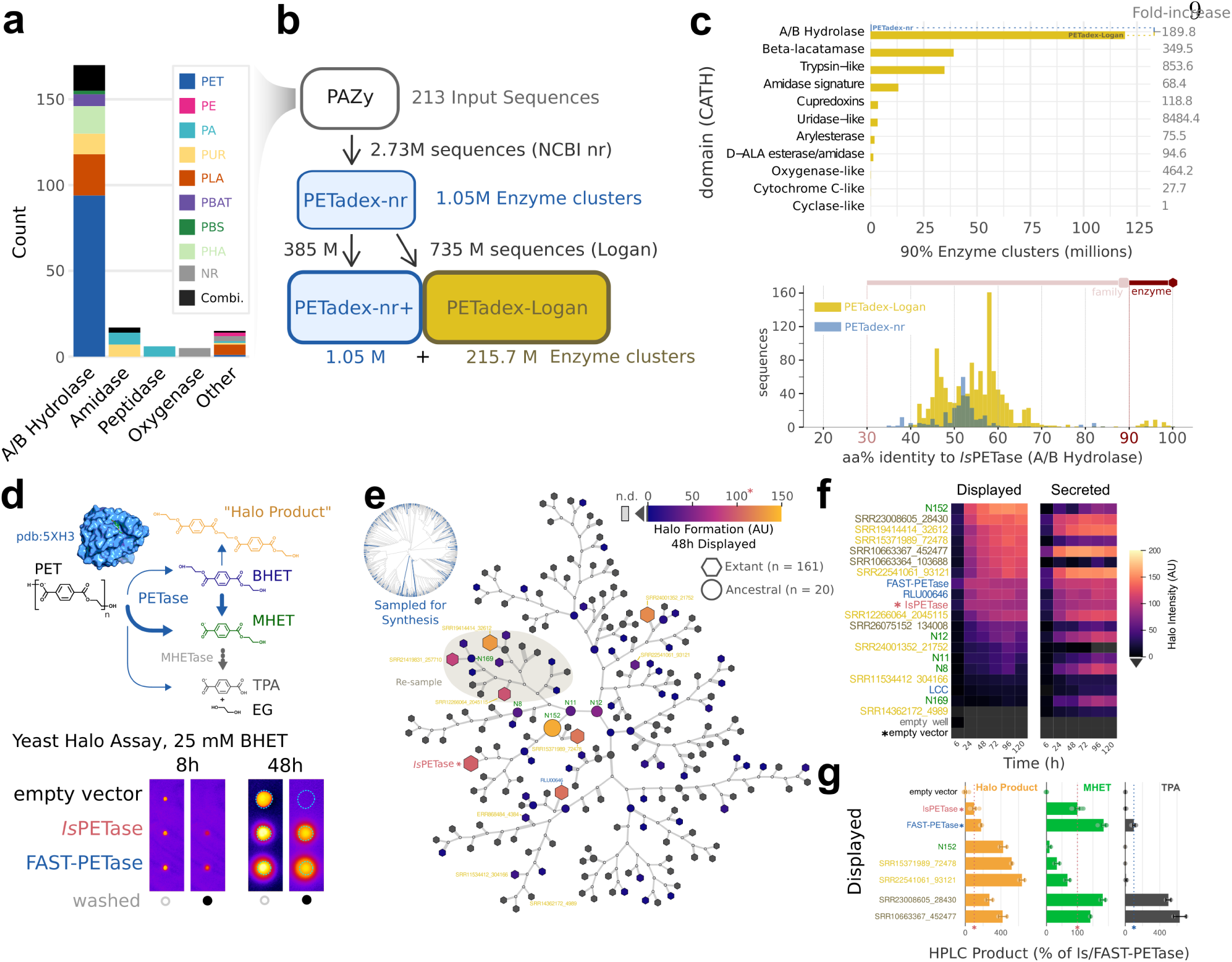
Discovering novel and efficacious plastic-active enzymes. **(a)** The domains and activity of the 213 experimentally validated plastic-active enzyme (PAZy) search query (Extended Data Fig. 2a) **(b)** Logan *PETadex* homology search returned 216.75 million PAZy-homologs after clustering at 90% amino acid identity (Extended Data Fig. 2b,c). **(c)** *PETadex* -Logan is a *>*200-fold expansion of candidate PAZy relative to NCBI nr across distinct PAZy CATH domains (see Methods). Histogram shows the distribution of *Is*PETase-aligned sequences, illustrating that Logan (yellow) uncovers more diversity across the detectable range of sequence identities relative to NCBI nr (blue). **(d)** The PETase reaction which underpins the high-throughput yeast-based halo assay. Yeast expressing either control (*Is*PETase) or candidate enzyme targeting the PET substrate BHET were grown on agar plates to create a white halo which is quantified as pixel intensity (shown as pseudocolored), before (open circle) and after washing (black circle) around the colony (cyan outline) (Extended Data Fig. 3a,b). High-performance liquid chromatography (HPLC) and mass spectroscopy suggest that the “halo product” is O,O*^′^*-(ethane-1,2-diyl) bis(oxy(2-hydroxyethyl)carbonyl)terephthalate (Extended Data Fig. 3e,f). **(e)** Phylogenetic tree of sampled candidate PAZy that were synthesized and experimentally screened. Nodes are colored based on 48 hour halo formation activity in surface-displayed expression. The gray-highlighted clade was re-sampled for additional sequences. **(f)** Heatmap of select enzyme halo formation activity over time, quantified in surface-display and secreted systems (Extended Data Fig. 4a). **(g)** Quantitative validation of candidate high-activity *PETadex* -Logan enzymes by HPLC. The bars show the percentage of product formed relative to the activity of *Is*PETase (halo product, MHET) or FAST-PETase (TPA). Logan enzymes demonstrate product formation exceeding that of *Is*PETase and FAST-PETase.

To expand this, we then queried these 1.05 million enzymes across Logan contigs, recovering 1.12 billion matching sequences. From these, 385 million sequences (34.3%) matched an nr enzyme at 90% identity. We clustered the remaining 735 million sequences (65.7%) into 215.7 million non-redundant enzyme homologs (Fig. 2b). Overall, Logan provided a 205-fold expansion of sequence diversity over NCBI nr, spanning multiple plastic-active domains, including a 190-fold expansion of A/B hydrolases, of which *Is*PETase is a member (Fig. 2c). This dataset, *PETadex*, represents the most comprehensive and diverse collection of candidate plastic active enzymes. To facilitate the development of global research solutions for plastic remediation, we are releasing the *PETadex* data freely, and without restriction (https://github.com/ababaian/petadex).

To test if *PETadex* candidate plastic-active enzymes contain catalytically active sequences, we developed a quantitative high-throughput enzyme screen using a PET subunit substrate, bis(2-hydroxyethyl) terephthalate (BHET). Candidate PETase enzymes were expressed as cell surface displayed or secreted constructs in baker’s yeast, *Saccharomyces cerevisiae*. PET conversion activity was measured via a colometric halo around yeast colonies, which corresponds to the formation of 2-hydroxyethyl terephthalate (MHET) or a higher molecular weight “halo product” (Fig. 2d, Extended Data Fig. 3).

As an initial screen, we selected full-length *Is*PETase or PAZy A/B hydrolase-like homologs (40% aaid). From 2,272 unique matches, we synthesized 161 randomly selected enzymes, and 21 ancestral reconstructions (AR). We screened these enzymes over six timepoints and in the two expression systems, which revealed 35/161 (22%) of the natural, and 8/21 (38%) of the AR sequences had plastic-activity (Fig. 2f, Extended Data Fig. 4a). The most active enzymes identified showed overall conversion comparable to *Is*PETase with varying preferences for MHET or halo product formation (Extended Data Fig. 4a). Inspection of the enzyme phylogeny revealed a clade enriched for halo-formation activity, which was re-sampled for an additional 13 enzymes. Resampling yielded 9/13 (69%) plastic-active enzymes. High-performance liquid chromatography (HPLC) assessment of BHET conversion activity of the two most active enzymes (SRR23008605 28430 and SRR10663367 452477) revealed that these enzymes exceeded *Is*PETase activity for MHET or halo product formation (Extended Data Fig. 4b). Interestingly, these two enzymes also produced the PET monomer product TPA at substantially higher rates than *Is*PETase and 4-fold more TPA than the engineered FAST-PETase (Fig. 2g).

Microplastics are projected to increase exponentially and biocatalysts are a means by which these environmental contaminants can be remediated. To address this we created *PETadex*, a free and unrestricted resource of candidate plastic-active enzymes, two orders of magnitude more expansive than previously available. Logan resources such as this enable the deep exploration of the evolutionary landscape of proteins including identifying candidate enzymes with higher application-specific activities, more complete product yields (TPA formation), or novel chemical functions (halo product formation).

### Characterization of Human Herpes Virus 6 reactivation in heterogeneous *ex vivo* cultures

Our recent work has demonstrated that retrospective assembly and quantification of viral nucleic acids at the petabase scale could reveal new associations between humans and viruses, including in clinical contexts [2],[22]. Specifically, comprehensive mining of Serratus [2] led to the discovery of Human Herpesvirus 6 (HHV-6) reactivation in chimeric antigen receptor (CAR) T cells [22], a finding that contributed to revised Food and Drug Administration guidelines requiring the screening of viral reactivation in allogeneic CAR T cells [23].

We hypothesized that Logan could further identify instances of viral reactivation in human cells and tissues where viral expression was not considered in the primary analyses. Among the 103 HHV-6 type B (HHV-6B) genes, we selected two transcripts U83 and U91 for sequence query based on high expression, short length (less than 1kb), and known gene function (Extended Data Fig. 6a; Methods). We queried these two transcripts using Logan-Search against 1,476,236 human RNA-sequencing datasets (Fig. 3a). Each query took less than five minutes to complete, resulting in 13 distinct BioProjects with *>*50% *k*-mer coverage across both HHV-6 transcripts (Fig. 3b; Methods), four of which had known HHV-6 expression. These four projects served as positive controls, which included CD4+ memory T cell cultures annotated by Serratus [24],[25] and CAR T products with previously characterized HHV-6 reactivation [22], [26]).

**Figure 3:**
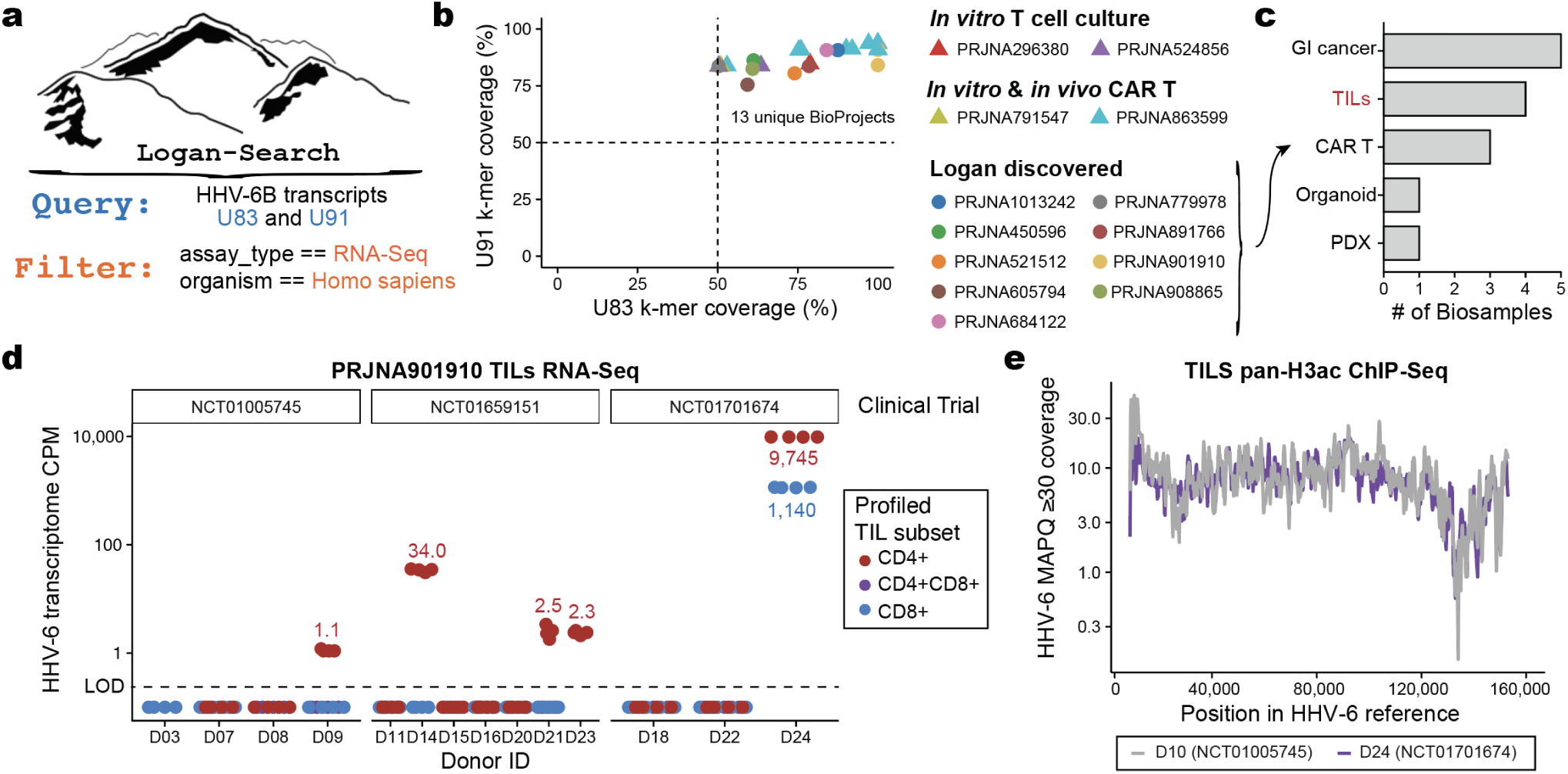
Identification and reactivation of HHV-6 in large-scale RNA-seq datasets. **(a)** Input query to Logan-Search for two abundant HHV-6 genes (U83 and U91) and filtering criteria for human RNA-seq BioSamples. **(b)** K-mer coverage analysis of 13 identified HHV-6–positive BioProjects, including 9 with no prior HHV-6 annotation (circles). Triangles indicating previously annotated datasets (Serratus and HHV6 paper). **(c)** Annotation of newly discovered HHV-6–bearing BioSamples, including gastrointestinal cancers, tumor-infiltrating lymphocytes (TILs), chimeric antigen receptor (CAR) T-cell products, organoids, and patient-derived xenografts (PDXs). **(d)** RNA-seq analyses of TIL cultures (PRJNA901910). Values indicate HHV-6 RNA abundance (counts per million, CPM) out of the full library, reflecting HHV-6 reactivation from cultured T cells. **(e)** ChIP-seq analyses of TIL cultures (PRJNA901909). Values reflect the HHV-6 DNA abundance for two donors in H3 acetylation chromatin, reflecting coverage across the full viral contig.

Supported by the recovery of these positive controls, we then considered the nine novel BioProjects. According to the BioSample meta-data, these samples comprised additional gastrointestinal tumors [27] and CAR T cells [22], consistent with previous characterizations in other settings (Fig. 3c). The novel CAR T BioSamples were from a study profiling infusion products and longitudinal profiles from 26 patients with B cell acute lymphoblastic leukemia receiving anti-CD19/CD22 CARs [28]. While this study focused predominantly on CAR-intrinsic gene expression changes associated with variable patient outcomes, Logan enables retrospective discovery of HHV-6 reactivation in these samples. Further, as this study was published after the completion of Serratus [2], HHV-6 reactivation in this cohort was not previously annotated or reported by the original authors. These results further supports our prior conclusions of HHV-6 reactivation occurring agnostic of disease or CAR target.

Next, we focused on the tumor-infiltrating lymphocytes (TILs) or organoid model annotated BioSamples. To the best of our knowledge, no prior reports of viral reactivation had been previously reported in these settings. However, since these systems involve extended cultures of heterogeneous mixtures that include CD4+ T cells, we reasoned that the Logan associations could reflect HHV-6 reactivation in *ex vivo* cell culture settings of T cells [22]. Indeed, in the lung organoid sample [29], we identified a population of HHV-6 super-expressor cells among the CD4+ proliferating T cells (Extended Data Fig. 6b), supporting our prior characterization of a rare cell state responsible for seeding lytic virus following reactivation [22]. The annotated HHV-6+ TIL samples were infusion products profiled from patients with metastatic melanoma treated from three clinical trials [30]. Analysis of the 16 donors profiled with RNA-seq demonstrated high-confidence HHV-6 detection in 5 infusion products, predominantly from the CD4+ sorted populations (Fig. 3d). At least one positive donor was observed in each trial, underscoring that HHV-6 reactivation in these adoptive cell therapies is a recurrent phenomenon.

As viral RNA accumulation coincides with increased viral DNA copy number [22], we examined additional profiles of clinical TIL products analyzed via chromatin immunoprecipitation and sequencing (ChIP-seq) for a pan-H3 acetylation modification that marks transcriptionally active chromatin. Among 19 donors spanning the same three clinical trials, we observed viral reactivation in samples from all three trials, including high HHV-6 expression from a donor (D10) where RNA-seq was not obtained (Fig. 3e; Extended Data Fig. 6c). Further analyses of viral single-nucleotide polymorphisms revealed 72 mutations specific to either donor, excluding the possibility of a common source of HHV-6 contamination during library preparation (Extended Data Fig. 6d). Across both modalities, our results suggest that HHV-6 reactivation in T cell therapies occurs independent of exogenous DNA and further implicates the rapid proliferation of T cells *ex vivo* as a critical signal underlying HHV-6 reactivation *in vitro*.

Taken together, our analyses shows that Logan effectively uncovers novel biological associations of viruses using existing human genomic profiles. In particular, this vignette reveals that latent HHV-6 can reactivate in rare proliferating CD4+ T cells from heterogeneous cell culture conditions, spanning from organoids to adoptive cell therapies with or without genetic engineering. As culture duration is a key determinant of viral reactivation in CAR T cells [22], our observation of HHV-6 reactivation in TIL therapies is consistent with the longer culture durations than widely-used autologous CAR T cell therapies. Our characterization of viral reactivation *ex vivo* motivates further work into gene editing and/or small molecule approaches that can mitigate reactivation to maximize the safety and efficacy of cell therapies [23]. More generally, our results motivate future work to monitor viral reactivation across current and future cell therapies using comprehensive genomics profiling and scalable analyses enabled by Logan.

### Expanding the Known Universe of Proteins, Plasmids, and Viral Elements

Next, we mined the 0.9 petabases of Logan’s assembled contigs to reveal order-of-magnitude expansions in the known diversity of proteins and mobile genetic elements. These planetary-scale deep homology searches were completed in as little as 11 hours, using cloud-deployed translated protein-to-nucleotide alignment.

#### Billions of diverse Logan proteins

Logan expands the known protein universe, with 109.4 billion proteins clustered into 3.0 billion non-redundant sequences at 90% amino acid identity and 90% alignment overlap (Fig. 4d). This represents over an order of magnitude greater set of protein diversity relative to large-scale commercial (BaseData [31]) or public (OMG [32], MGnify [13], BFD [33]) metagenomic resources, and a nearly 30-fold increase over UniRef50.

**Figure 4:**
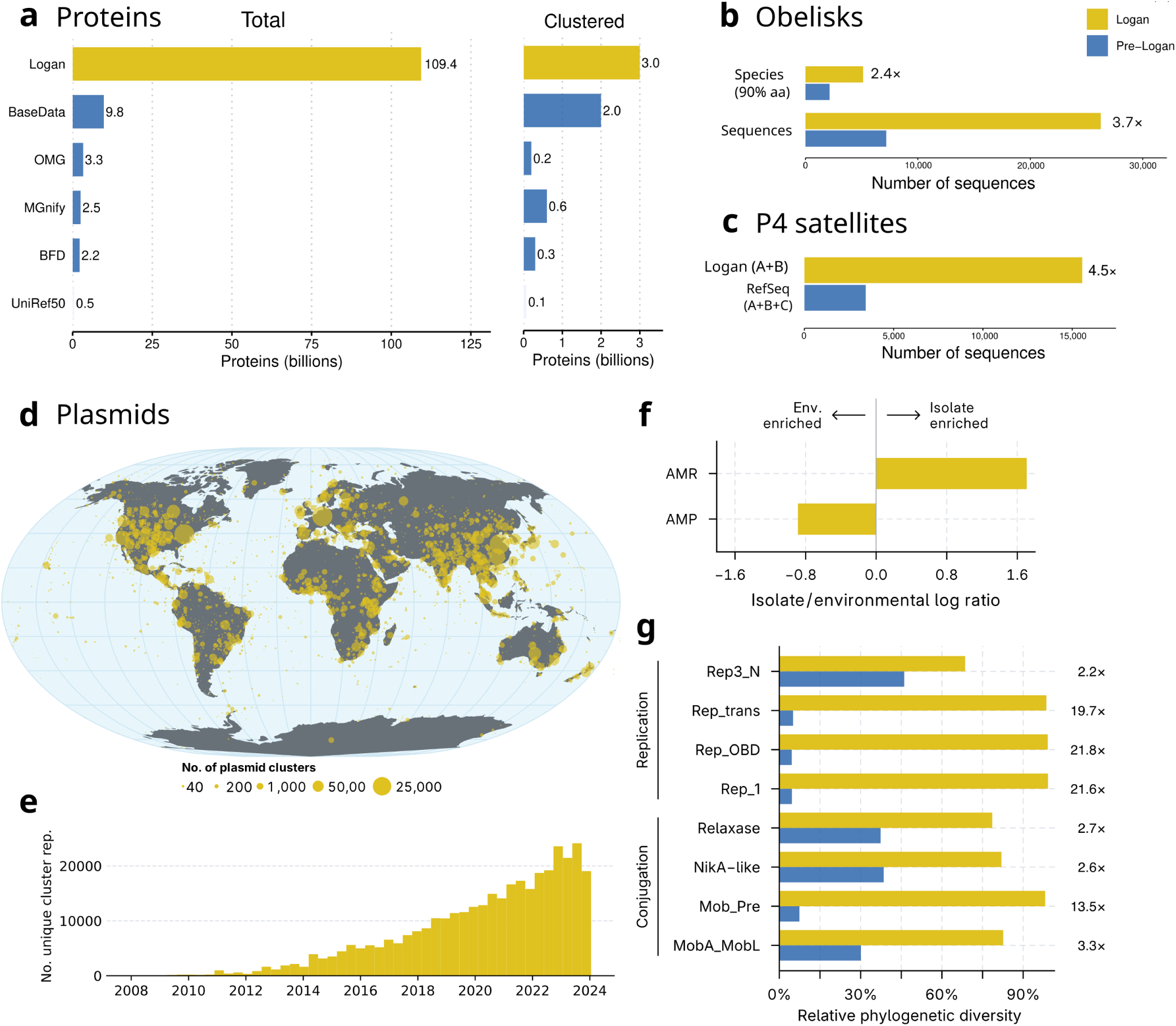
Expanding the Known Universe of Proteins, Plasmids, and Viral Elements. **(a)** Bar plot labeled “Total” shows the total number of proteins extracted from Logan contigs, compared to other databases, and bar plot labeled “Clustered” shows the same set of proteins but clustered at 50% identity. **(b)** Expansion of Obelisks requiring circular contigs with full-length Oblin-1 proteins, identified in Logan (yellow), relative to the initial petabase-scale search [9] (blue). Total sequences and species (clustered centroids of Oblin-1) are shown. **(c)** Number of P4 satellites found in Logan contigs (types A+B) compared to those in RefSeq (types A+B+C). Types A, B, C refer to the number of core components detected by SatelliteFinder (Methods): A = all 7, B = 6 out of 7, C = 5 out of 7. **(d)** Geographical distribution of plasmids detected over 2,095,914 samples with geolocation data. Circle areas are proportional to the number of distinct plasmid clusters per region. For visual clarity, spatially close samples were grouped using DBSCAN, and circles are placed at the coordinates of the corresponding cluster medoids. The map uses the Loximuthal projection. **(e)** Number of distinct plasmid cluster representatives detected over time. Counts (y-axis) are shown in 120-day intervals (x-axis). **(f)** Comparison of accessory gene repertoires in plasmids from environmental samples vs. cultured isolates. Plasmids from isolates are comparatively enriched for antimicrobial resistance (AMR) genes, whereas plasmids from environmental sources are enriched for antimicrobial peptides (AMPs). Functional enrichment (x-axis) was quantified as the ratio of gene density (genes per megabase) for a given function (AMR or AMP) between the two plasmid groups. **(g)** Relative Faith’s phylogenetic diversity of selected replicases and relaxases encoded by plasmids identified in Logan (yellow) and those retrieved from PLSDB (blue). The phylogenetic diversity fold change, indicated by numbers to the right of the bars, represents the ratio of the diversity across all plasmids (Logan and PLSDB combined) to the diversity in PLSDB plasmids alone.

This expanded diversity provides an invaluable resource for downstream applications. In a case study of 100 viral proteins, sensitive searches against a clustering of Logan proteins at 50% amino acid identity produced substantially more diverse multiple sequence alignments compared to searching against a standard database [33]. We observed an approximately 2-fold increase in the Number of Effective Sequences (Neff: 2.19 to 4.89; Extended Data Fig.5B, left panel). Doubling of Neff values translates into improved protein structure predictions, with mean predicted Local Distance Difference Test (pLDDT) scores rising from 46.7 (“very low”) to 88.6 (“high”), and 90 out of 100 proteins modeled at high quality (Extended Data Fig.5B, right panel), highlighting the impact of our expanded protein clustering on improving AI-structure prediction accuracy.

#### Obelisks

Obelisks are a newly discovered clade of viroid-like agents. Initially two complete, circular species were identified as persistent colonists in human gut metatranscriptomes, then a petabase-scale SRA search expanded this to 2,152 species genomes (defined here as Oblin-1 90% aaid clusters) [9]. With Logan we detected an additional 2,964 Obelisk species, a 2.4-fold increase. Likewise, the total Obelisk count increased 3.7-fold, to 26,263 sequences (Fig. 4b). The expanded Oblin-1 proteins have no homologs in NCBI nr, and no homologs known outside of Obelisks. This expanded dataset should accelerate research to uncover the role of these mysterious elements.

#### P4 Satellites

We then searched for P4-like satellites [34], mobile elements that hijack bacteriophages. P4-like satellites were recently found to be numerous in enterobacterial genomes, where they provide the host with anti-phage functions. We observed a 4.5-fold expansion of the number of these satellites (Fig. 4c), including elements that are unrelated with previous sub-families (Extended Data Fig. 8c). The pan-genome of these elements is thus doubled in relation to RefSeq (Extended Data Fig. 8c). These newly uncovered P4 elements may thus encode many novel anti-phage functions of ecological and biotechnological relevance. These searches illustrate how to leverage Logan to mine for complex genetic elements with multiple core genes.

#### Plasmids

Plasmids are mobile genetic elements of Bacteria and Archaea that play a critical role in horizontal gene transfer, driving processes such as the spread of antibiotic resistance genes. Given their ecological and clinical importance, systematically characterizing plasmid diversity can provide valuable insights into global gene transfer patterns. We extracted all circular contigs from Logan assemblies and identified 468,614 putatively complete plasmids (264,160 unique sequences) across 195,347 metagenome, 57,148 bacterial, and 12 archaeal isolate accessions. To evaluate the global distribution of these plasmids, including in samples where they were not fully assembled as circular contigs, we first reduced redundancy in the dataset by grouping highly similar sequences into 60,331 clusters, and then mapped representative sequences from each cluster to all Logan contigs. This approach identified plasmids in 2,095,914 samples over the globe (Fig. 4d), highlighting their widespread distribution. Moreover, the number of distinct detectable plasmids has steadily increased over time (Fig. 4e).

Next, we investigated the origins underlying this extensive plasmid diversity by analyzing the composition of plasmid clusters. We found that 92.5% comprised only metagenomic sequences, while just 3.6% consisted exclusively of plasmids from cultured organisms. Despite this predominance of environmental sequences, only 17.5% of metagenomic plasmids could be assigned to known replicon families compared to 78.3% from isolates, highlighting that the plasmid diversity in natural environments remains largely uncharacterized. Environmental plasmids were depleted of known antimicrobial resistance (AMR) genes and encoded more antimicrobial peptides (AMPs) relative to plasmids from isolates (Fig. 4f), reflecting distinct accessory gene repertoires and underscoring the value of assessing plasmid diversity across diverse sample types, as enabled by Logan.

To quantify the extent of plasmid diversity uncovered by surveying Logan assemblies, we measured the phylogenetic diversity of selected replicase and relaxase proteins from these plasmids and compared it with complete plasmid genomes from established databases [35, 36]. Plasmids identified from Logan assemblies expanded the phylogenetic diversity up to 21.8-fold compared to PLSDB (Fig. 4g) and up to 7.0-fold compared to IMG/PR (Data Availability, Plasmid PD Table). Overall, these results demonstrate that mining Logan assemblies reveals a vast and previously undiscovered diversity of genetic elements not captured by other genomic data resources.

#### Antimicrobial resistance

Logan also enables the global-scale analysis of antimicrobial resistance (AMR) across all publicly available sequencing data. By aligning all Logan contigs to the CARD database, we identified 7.9 million AMR-positive (AMR+) SRA accessions and 13,000 AMR+ plasmids (Extended Data Fig. 7a). AMR genes are enriched in metagenomes across the SRA, whereas plasmids show the opposite pattern, driven by the large fraction of bacterial isolates in plasmid datasets (Extended Data Fig. 7b). We observe an enrichment of human and livestock metagenomic samples in AMR-positive datasets (Extended Data Fig. 7c), both in SRA accessions and plasmids. Geographically, AMR+ metagenomes are broadly distributed, and their discovery has increased steadily over the past two decades based on collection dates (Extended Data Fig. 7d-e). AMR gene content varies across metagenome categories, with wastewater, livestock, and human metagenomes showing the highest enrichment in AMR gene counts per accession (Extended Data Fig. 7f-g). As sequencing databases continue to grow, full-scale sequence indexes can be re-purposed as an AMR-surveillance network.

## 2 Discussion

The exponential growth of sequence databases has created a paradox: humanity is generating more genomics data than ever before, which is making the data increasingly inaccessible due to computational barriers. Logan resolves this paradox through transforming raw sequencing data into accessible and searchable resources which enable systematic analysis of global sequence diversity, and offers a technical framework by which analysis capacity can continue to scale alongside database growth.

Earth’s genetic diversity is a heritage of humanity [37]. To bulwark against a trend of commercializing public scientific data, we release all Logan data into the public-domain, and emphasize the continued need for community development of free and unrestricted data commons (https://registry.opendata.aws/pasteur-logan/). For decades, BLAST democratized sequence comparison at the gigabase scale, and it transformed biology research. Logan brings such a capacity to the petabase-era, enabling analogous discoveries across orders of magnitude larger datasets. Like the original NCBI web server that made BLAST a ubiquitous tool, Logan-Search’s web-interface (https://logan-search.org/) ensures that this resource is practically available for the research community.

Logan-Search enables researchers to rapidly test hypotheses across the breadth of public sequencing data, uncovering unexpected connections that were previously hidden in plain sight, with direct applications to biotechnology and planetary health. The discovery of HHV-6 reactivation in therapeutic CAR T-cells illustrated how large-scale sequence search could inform clinically relevant insights [22]. Here, we generalize such capability and extend this discovery to characterize latent viral reactivation tumorinfiltrating lymphocytes. Logan enables researchers to query the sampled biosphere for specific functions, such as identifying novel plastic-degrading enzymes that outperform engineered variants. It allows us to directly sample from nature’s vast parallel evolutionary experiment.

The massive expansion of plastic-active enzyme homologs makes it immediately obvious that Logan enables the free and unprecedented capacity to screen for variants and novel versions of proteins and genes for biotechnology. This includes but is not limited to, for example, identifying novel viral vectors, or receptors/effectors with altered tropism; biosynthetic gene clusters for the production of antibiotics or natural products; efficient or process-optimized industrial enzymes such as proteases, amylases, or cellulases; or biotechnology enzymes such as Cas, reverse-transcriptases, or polymerases. Moreover, these public datasets are a rich resource with obvious applications as training data for a next-generation of machine learning and artificial intelligence models.

The ability to efficiently analyze all public sequence data arrives at a critical moment for biodiversity research. Current sampling of Earth’s genetic diversity shows clear geographic and taxonomic biases, evident in SRA metadata. As climate change and habitat loss accelerate species extinction, systematic sequence analysis becomes essential not only for documenting disappearing diversity but for understanding the genetic basis of adaptation and resilience. Logan provides the technical foundations, while highlighting the urgent need for broader and more representative sampling of Earth’s biosphere.

## 3 Methods

### 3.1 Performance-optimized and cloud-accelerated genome assembly of all SRA accessions

We designed a cloud architecture to perform SRA-wide genome assembly efficiently and in parallel. Figure 1b describes the workflow. Briefly, a Docker container processes each SRA accession independently: 1) raw reads of an accession are downloaded from a cloud mirror of the SRA, 2) a conservative assembly (unitigs) of the accession is made from the reads, 3) a consensus assembly (contigs) is made from the unitigs, and 4) both the unitigs and contigs are compressed and uploaded to a public repository.

We executed this system at SRA-scale using cloud resources. Containers are executed in parallel over tens of thousands of cloud computers through a container orchestration system, and a set of dashboards were deployed to monitor the execution. Extended Data Fig. 1 shows some key statistics of the execution. We optimized the system to use as many CPU cores in parallel as possible, as opposed to running it at smaller scale over a longer period of time, to take advantage of lower computation costs and higher availability of cloud instances during night time.

Using this system we performed genome assembly over the entire SRA and report results for each accession in two forms: unitigs (non-branching paths of the reads de Bruijn graph, here *k* = 31) and contigs (non-branching paths of the de Bruijn graph after simplification steps). The rationale for this dual set of results is to allows researchers to choose between a near-lossless but fragmented representation (unitigs) and a more lossy but more contiguous one (contigs). Unitigs are shorter sequences that preserve almost all the genetic variation from the original reads, including alternate alleles involving single nucleotide or indel variants. Unitigs are ideal for sensitive, *k*-mer-based searches where finding even minor variants is critical. Contigs, on the other hand, are the more contiguous, consensus assemblies. To obtain contigs, the assembler extends the unitigs by collapsing biological variations and removing any remaining putative technical sequencing error. It makes a “best guess” of the errors to remove and major allele to collapse into a single, representative path. Contigs are optimized for downstream applications like protein prediction and gene identification, where sequence contiguity is more important than preserving minor variants.

#### 3.1.1 Input data

We selected all public samples from the Sequence Read Archive on December 10th, 2023, with read length above 31 bp. Accessions with shorter reads than 31 bp would yield no usable *k*-mers in the downstream assembly step. The list of accessions was obtained from the NCBI SRA metadata table, using Amazon Web Services (AWS) Athena with SQL query filter: WHERE consent = ‘public’ and avgspotlen >= 31. This resulted in 27,764,168 accessions totalling 50,304,659,857 bases in reads. In the rest of this manuscript we refer to this dataset as ‘the SRA’, although the current-day SRA has since been updated with new samples.

#### 3.1.2 Assembly tools

Unitigs were constructed using a modified version of Cuttlefish2 [38] (commit 9401ef5 of forked repository github.com/rchikhi/cuttlefish), augmented to record approximate mean *k*-mer abundance per unitig. We also modified the *k*-mer counting method KMC3 [39] integrated inside of Cuttlefish2 to stream SRA files directly through a piped call to fasterq-dump with parameters --seq-defline ‘>’ --fasta-unsorted --stdout, avoiding a prior decompression step to disk, and discarding on the fly FASTQ headers and quality values.

The original version of Cuttlefish2 did not record any abundance value. We modified Cuttlefish2 to record abundances values per *k*-mer during construction, then to report the average abundance over all *k*-mers of a unitig. However, the reported per-*k*-mer abundances were approximated with two heuristics: 1) due to a technicality during graph construction, abundances of *k* + 1-mers were recorded, hence the abundance of each *k*-mer was obtained by summing the abundances of all the *k* + 1-mers it appeared in, then dividing the sum by two. 2) To save memory during graph construction, abundances were stored in a 8 bits encoding scheme that maintains an error of not more than 5%, with a maximum abundance value of 50,000. To remove some of the likely sequencing errors, *k*-mers seen only once in an accession were discarded from unitigs.

Unitigs were given as input to Minia3 [40] (commit 71484e8 of github.com/GATB/minia), which performs de Bruijn graph simplifications and error-correction following closely the heuristics designed by the SPAdes assembler [41], a tool well-established for its high accuracy and low rate of misassemblies in metagenomic contexts. We expect Minia3 contigs to share a similar accuracy profile, since the modifications focused primarily on performance optimization and memory frugality, rather than altering the fundamental algorithms responsible for assembly quality. Minia3’s accuracy was independently validated in [42, 43, 44]. The resulting contigs that are longer than 150 bp and are connected to at least one other contig were reported. For both unitigs and contigs, the FASTA headers contain link information (in the format of BCALM2 [45] output) that enable to reconstruct the assembly graph in GFA format. In the unitigs and contigs file of an accessions all 31-mers are distinct, by construction.

These tools were selected for their memory and running time frugality. In addition, Minia3 was chosen for also its conservative approach to graph simplification and hence higher retention of sequence complexity. A comparison with two other state-of-the-art assemblers (Penguin [46], rnaviralSPAdes [47]) is provided in Extended Data Fig. 1, showing that substantial cloud computing costs were saved by this pipeline.

To compress unitigs and contigs, we developed and applied a novel block variant of the Zstandard algorithm [48] (https://github.com/asl/f2sz). The compressor creates FASTA-aligned blocks allowing for faster random access to any subset of contigs. It remains compatible with the ubiquitous zstd -d and zstdcat decompression command line tools.

#### 3.1.3 Cloud infrastructure

For SRA-scale assembly we have a set up a cloud infrastructure on Amazon Web Services (AWS), to perform assembly of each SRA accession in a fully parallel fashion. In brief, the infrastructure is based on a Docker container executing a set of Python scripts responsible for calling child programs for unitig and contig assembly and validation. The AWS Batch execution system handles scheduling of containers across a pool of cloud computers (AWS EC2 instances). Each container is executed independently for each SRA accession. Each instance is equipped with temporary network storage (EBS). The number of CPUs, RAM, and storage for each job were set according to size of input reads measured in megabases. The Batch instances pool was set to the c6g and c7g families (AWS Graviton-based instances), with sizes 4xlarge and above to target larger instances and thus limit the number of instance creation API calls.

Executions were monitored primarily through live dashboards on AWS CloudWatch and Batch services, as well as a DynamoDB database recording runtime and assembly statistics for each accession. Global statistics such as number of processed accessions and total size of raw data assembled were recorded in real-time by sending messages from each container to a global partitioned database. Summary metrics were aggregated, enabling to monitor computation speed during execution and potentially stop all jobs should metrics fall behind projected estimates.

To limit the number of simultaneous queries to NCBI servers, two mechanisms were implemented: (1) raw .sra files were directly downloaded from the AWS Registry of Open Data cloud mirror of the NCBI SRA to a cloud instance in the same data center (us-east-1), and (2) special .sra files containing alignments to RefSeq were handled by downloading references from a S3 mirror of RefSeq. Aligned .sra file containing references from the NCBI WGS database were discarded in the later runs, but some were processed in the earlier runs before we identified that they incurred (rate-limited) queries to NCBI servers.

Results (unitigs, contigs) have been deposited in a public repository. Detailed instructions to download the data are provided here: https://github.com/IndexThePlanet/Logan.

While the cloud infrastructure used to construct Logan is solely intended for internal usage due to its high execution costs, its source code is publicly available at https://gitlab.pasteur.fr/rchikhi_pasteur/erc-unitigs-prod/.

#### 3.1.4 Assembly results

In total 27.3 million accessions were assembled into unitigs, representing 96% of the SRA in size as of December 2023. Some accessions resulted in too many unitigs to fit the assembly graph in memory, hence were not further assembled into contigs at this time. 26.8 million accessions were assembled from unitigs to contigs, representing 88% of the SRA in size. The total cloud computation time (unitigs and contigs) was around 30 million CPU hours (1).

Assembly contiguity statistics for the Logan contigs are reported in Extended Data Fig. 1. Contigs for Whole-Genome Sequencing/Amplification (WGS/WGA) accessions are generally longer than those of RNA-Seq accessions or other sequencing types, as expected by the longer sequenced molecules. Note that non-circular contigs shorter than 150 bp that are isolated nodes in the assembly graph were discarded by the assembler as they were more likely to be artifacts than actual biological material.

Across Logan, standard assembly metrics were computed using seqkit [49] and stored into a database (number of unitigs, contigs, N50 values, total length of assemblies, longest assembled sequence per accession). In addition, FASTA file sizes before and after Zstandard compression were also recorded. All these statistics were stored on a AWS DynamoDB database then exported to a public repository (S3 bucket) in the Parquet format (https://github.com/IndexThePlanet/Logan/blob/main/Stats-v1.md), enabling users to link this database with other NCBI SRA databases such as STAT [5] or SRA metadata.

### 3.2 Shallow and deep homology search in Logan contigs

For translated-protein searches over all Logan contigs, DIAMOND2 v2.1.9 [11] was run with parameters -b 0.4 --masking 0 -s 1 --sensitive to balance speed and sensitivity. The query sequence(s) were provided as indexed references to DIAMOND2, and Logan contigs were streamed accession by accession as queries. For nucleotide searches over all Logan contigs, minimap2 v2.28 [50] was run with parameters -x sr --sam-hit-only -a to return all contig sequences and their alignment, optimizing for short matches. A custom cloud pipeline using AWS Batch was set up to perform those searches, which each take approximately 11 hours on 60,000 vCPUs. The pipeline and infrastructure source code is available at https://gitlab.pasteur.fr/rchikhi_pasteur/logan-analysis.

### 3.3 SRA-wide public search engine

We developed Logan-Search, a publicly available search engine able to approximately locate a queried DNA sequence among 23.4 million accessions. Logan-Search performs *k*-mer-based queries. A *k*-mer is a word of length *k* (*k* = 31 is used in this context). Within minutes, Logan-Search identifies the accessions to which each *k*-mer from the queried sequence is associated. This enables us to provide a similarity metric between the query and each indexed accession.

Constructing the Logan-Search engine required to index all *k*-mers from Logan unitigs, in order to offer a way to instantaneously detect to which sample a *k*-mer belongs to. We built this index using kmtricks [51] (kmtricks-logan tag at github.com/tlemane/kmtricks) and kmindex [52] (v0.5.3). Computations were performed on Microsoft Azure cloud platform, using an Azure Batch Pool consisting of 625 virtual machines (VMs) of type Standard D32d v5. The workload was divided into approximately 45,000 tasks used to construct partial indexes, which were subsequently merged to produce the final index. A small fraction of tasks requiring larger memory capacity were executed on Standard D96d v5 instances. In total, the computation took around 10 days to complete.

In total, approximately 2 × 10^15^ *k*-mers were indexed. The accessions were separated into groups on the basis of their library source (e.g. genomic, transcriptomic, metagenomic, metatranscriptomic, etc…) and of their superkingdom phylogeny classification, obtained from the NCBI STAT database [5]. Exceptions were made for human and mouse accessions, which were classified separately. The kmindex tool builds Bloom Filters [53], with a non-null false positive rate, tuned to be approximately 0.005% by setting parameters adapted to the size of each indexed dataset and by using the Findere algorithm [54]. The overall size of the index is approximately one petabyte, stored on disk. At query time, only specific target sub-parts of the index are mapped on RAM. For a thousand basepair sequence query, results are obtained in approximately 6 minutes, using 12 small 4-vCPUs virtual machines.

The kmindex tool retrieves accessions containing sequences similar to a query, based on the percentage of *k*-mers from the query existing in the accession. On top of those results, we developed a visualization interface based on kmviz (https://github.com/tlemane/kmviz) that offers several features:

- All metadata from SRA associated to each accession is made available. The interface enables researchers to visually retrieve those metadata: geographic localization of accessions on a worldwide map, and drawing highly tunable plots from discrete or textual attributes.
- A summary of the query is generated using a large language model (GPT-4o) [55], based on the SRA metadata associated to top hits, allowing users to quickly assess potentially relevant contextual information, such as organism or location.
- Logan-Search natively returns a list of accessions that contain the query sequence, but it does not identify the specific sequence within each accession that matches the query. To fill this gap, a microservice based on the back to sequences tool [56] is used to retrieve, on a per-accession basis, the contig or unitig sequences that match the query. Additionally, a BLASTn [57] alignment using default parameters is performed between the query and the extracted contigs or unitigs; it completes instantly as both aligned sequences are typically kilobase-sized.

### 3.4 Pilot plastic active enzyme search

In a pilot experiment to assess the feasibility of a plastic-active enzyme search, we retrieved 102 publicly available, experimentally validated, polyethylene terephthalate (PET)-degrading A/B hydrolases from the PAZy database (sequence accessed Aug 26, 2024 from GenBank or PDB accessions [18]). These reference sequences were supplemented with 12 previously computationally identified PETases and 74 MGnify sequences with predicted structures similar to *Is*PETase, found with Foldseek [58, 59, 13]. To reduce redundancy, these 188 enzymes were clustered at 90% aaid (‘USEARCH v11.0.667 i86linux32‘, -cluster fast -id 0.90), resulting in 153 representative sequences [60].

We focused on 56 sequences related to *Is*PETase by at least 30% aaid (USEARCH v11.0.667 i86linux32, -cluster fast -id 0.30) [60]. We filtered Logan retrieved contigs to “*Is*PETase-like” sequences with high identity and confidence, resulting in 230,804 hits (‘DIAMOND2 blastx v2.1.9‘ as above; e-value *<* 1*e^−^*^8^ and aaid *>* 40%) [11].

As we were interested in potentially active enzymes with intact catalytic cores, we generated a multiple structure alignment of *Is*PETase-like reference sequences. Structures were predicted using AlphaFold3 and aligned with Muscle3D (v5.1.linux64, -align) [61, 62]. The amino acids of the subalignment containing the conserved core were extracted manually and were used to generate a custom HMM (‘HMMER v3.4‘; esl-reformat stockholm, hmmbuild) [63]. Next, we identified stop-stop ORFs (‘EMBOSS v6.6.0.0‘; getorf) and filtered for those with at least 90% coverage of the core HMM (‘HMMER v3.4‘; hmmsearch), resulting in 2,272 unique sequences with a maximum e-value of 3.8*e^−^*^43^.

To infer the evolutionary relationships of these Logan hits, we clustered them at 95% aaid (853 representative sequences), and used IQ-TREE2 with 1000 bootstrap iterations to generate a tree (v2.4.0, -b 1000) [64]. Ancestral reconstruction of these sequences was performed with GRASP command-line [65], resulting in 175 ancestral sequences. For initial synthesis and activity testing, 161 candidate sequences from Logan were manually chosen over the tree to encompass an even sequence diversity. Twenty-one ancestrally reconstructed sequences were included, as well as six distantly related sequences from MGnify as expected negative controls, and 11 known PETases as positive controls, amounting to 199 sequences in total.

After the PETase activity was measured for the initial round of synthesized sequences (see Section 3.8) the phylogenetic tree was re-sampled in areas of relatively enriched PETase activity through manual identification of closely related and sister sequences (n = 13) for additional *in-vitro* screening.

### 3.5 Plastic active enzyme homolog expansion

As the initial search seed, we used the PAZy database of experimentally validated plastic-active enzymes (Accessed Dec 19, 2024) [18]. All publicly available PAZy sequences were retrieved from GenBank by accession (213/245, 86.94%). To generate a PAZy phylogenetic network (Extended Data Fig. 2a), we performed all-vs-all alignment (usearch v11.0.667) between sequences (213 nodes), and retained significant alignments (3,195 edges, *≥*30% amino acid identity and e-value *<* 1*e^−^*^5^). The sequences clustered into 42 graph components, of 11 structurally distinct protein folds (CATHdb [21]), with minimum-spanning tree edges shown. The Alpha/Beta Hydrolase protein family are the most represented, with 170/213 (79.8%) of the sequences and 28/42 (66.7%) of the components.

Notably, Alpha/Beta hydrolases, amidases, beta-lactamase, and arylesterase protein-fold families have reported activity for more than one plastic substrate, and 16 individual sequences have reported activity for more than one plastic substrate. Thus plastic degrading activity is a polyphyletic trait, and suggests it may be an incidental, or off-target function of these enzymes, which broadly function to hydrolyze organic molecules such as lipids, esters, or carbohydrates. This suggests these enzymes and their homologs may contain additional plastic substrate activity.

To identify PAZy homologs, we queried the sequences into the NCBI non-redundant protein database (‘nr‘, accessed: 2024-12-27) [10] using DIAMOND2 (--very-sensitive) [11] which retrieved 5,593,290 unique sequences (e-value *<* 1*e^−^*^5^). Of these, 2,735,790 sequences contained a HMM match (‘hmmscan‘; *<* 1*e^−^*^5^ and 95% model coverage) against a PAZy protein domain (Pfam models: PF00082, PF00089, PF00144, PF00561, PF01083, PF01425, PF01522, PF01674, PF01738, PF02983, PF03403, PF03576, PF06850, PF07224, PF07732, PF09995, PF10503, PF10605, PF12146, PF12695, PF13472, PF20434, PF21419, PF24708). Incidentally, we did not include a model for Cyclase-like domains, which resulted in zero novel sequences beig called.

Next, we queried these centroids against our newly created Logan assemblage (26.8 million datasets, 50.0 Pbp, 11.5 hours), as per Section 3.2, and retrieved an additional 6.5 billion hits, (DIAMOND, evalue *<* 1*e^−^*^8^), which were filtered as above.

### 3.6 Yeast strains and PETase expression vectors

All yeast strains generated in this study are derivatives of DHY213 [*MAT* **a** *CAT5(91M) SAL1 MIP1(661T) HAP1 MKT1(30G) RME1(INS-308A) TAO3(1493Q) leu2*Δ*0 his3*Δ*1 ura3*Δ*0 met15*Δ*0*], a modified version of BY4741 [66] with increased sporulation efficiency, mitochondrial stability and efficient biosynthetic gene expression [67].

PETase encoding genes were expressed as surface displayed or secreted constructs from episomal plasmids derived from the pJC170 backbone [68] (gift from Jef Boeke) modified to contain an additional marker (*natMX*) and the surface display or secretion expression cassettes. For surface display, PETase encoding genes were fused with the Ccw12 secretion signal (amino acid 1 to 19) on their 5’ ends and to a flexible [AGSAGSAAGSG] linker, a Myc tag and the remainder of the Ccw12 amino sequence (amino 22 to 133) on their 3’ ends. For secretion, the architecture of the construct was the same as for surface display but without the cell wall anchoring domain of Ccw12 (amino 22 to 133). Finally, the empty vectors additionally contain a placeholder sequence with two inverted BsaI restriction sites between the Ccw12 secretion signal and the linker sequence in order to facilitate backbone cleavage for recombination or golden gate cloning. The sequence for these plasmids (pRLK152 for surface display; pRLK153 for secretion) is provided in the supplemental material.

### 3.7 Synthetic DNA cloning

PETase encoding genes were synthesized by Twist Biosciences (USA) and contained the standard Twist adapters and 40 base pairs of homology on the 5’ (5’ CGCTTCTATCGCCGCTGTCGCAGCTGTCGCTTCTGCCGCA) and 3’ (5’ GCGGGTTCTGCTGGTTCTGCTGCTGGTTCTGGTGAATTTG) ends to facilitate *in vivo* recombination in yeast. Between 10 and 20 ng of synthetic DNA was transformed in yeast using the standard lithium acetate method along with 25-50 ng of plasmid backbone (pRLK152 or pRLK153). Yeast transformants were selected on synthetic medium containing clonNAT antibiotic and lacking uracil (SD/MSG-ura+ clonNAT: 1.7 g/L yeast nitrogen base without amino acids without ammonium sulfate, 1 g/L monosodium glutamate, 20 g/L dextrose, 20 g/L agar, 100 µg/mL clonNAT). The pool of transformants for each construct was maintained as a single colony on a colony array.

### 3.8 BHET halo assay

To prepare assay plates, a 1 M BHET (CAS# 959-26-2, Sigma-Aldrich) solution was prepared by diluting BHET flakes into 100% DMSO and heating slightly until complete dissolution. The BHET solution was then added to YPD medium (10 g/L yeast extract, 20 g/L peptone, 20 g/L dextrose, 20 g/L agar, 100 µg/mL clonNAT) prior to pouring Omnitray plates (ThermoFisher). Plates were kept at room temperature to prevent BHET recrystallization.

Yeast strains expressing surface displayed or secreted *Is*PETase were pinned in 384-array format (4 colonies per construct) onto YPD+BHET plates using a colony pinning robot (Singer Instruments, United Kingdom) and incubated at 30°C for up to 5 days. After incubation, yeast colonies were washed off the plates using water and a cell spreader tool to reveal a white halo or a clearing (loss of opacity). Each plate was imaged before robotic pinning as well as before and after colony washing (spImager, SP Robotics Inc, Canada).

BHET halo measurement was implemented in R as follows: First colonies were identified using the gitter package and the resulting colony mask was applied to the plate image before pinning (background image) and after colony washing (halo image) to extract pixel coordinates corresponding to each colony area in the image. Pixel intensity was extracted from each image using the imager package and the median pixel intensity for each colony area was then determined in the halo image and the median pixel intensity for the same colony area in the background image was subtracted. To normalize pixel intensity values across plates, the average median pixel intensity obtained across 4 replicate colonies containing the empty surface display or secretion plasmids present on each plate was subtracted from all colony median pixel intensity values on the given plate.

### 3.9 High performance liquid chromatography

Isogenic clones were isolated from each colony with activity in the BHET halo plate assay and grown to saturation in YPD containing 100 µg/mL cloNAT at 30°C. The saturated culture was then diluted 1000-fold in YPD+cloNAT and grown for 24 hours at 30°C. 95 µl of culture was transferred into a 96-well plate prior to adding 5 µl of 500 mM or 250 mM BHET in 100 % DMSO. After 17 hours of incubation at 30°C, an aliquot of each reaction was diluted 10 or 20 times in 100% DMSO for reactions in 12.5 and 25 mM BHET respectively, centrifugated for 2 minutes at 3000 rpm and the supernatant was stored at -20°C.

Supernatants were fractionated on reversed-phase HPLC using an HP1050 system (HP/Agilent, USA) mounted with a Zorbax SB-C8 column (4.6 x 150 mm, 5 µm). The column was maintained at 22-24°C. Analytes were eluted over 37 minutes with 0.1% formic acid in water (aqueous solvent) and 0.1% formic acid in acetonitrile (organic solvent) using the following gradients: 1 to 5% organic (vol/vol) over 20 minutes at 0.8 ml/min, 5 to 52.5% organic (vol/vol) over 14 minutes at 0.8 ml/min, 52.5% to 100% organic (vol/vol) and 0.8 to 3.0 ml/min over 0.2 min, 100% organic (vol/vol) for 0.8 min at 3.0 ml/min, 100% to 1% organic (vol/vol) and 3.0 to 0.8 ml/min over 1.0 min, and 1% organic (vol/vol) for 0.2 minutes. Detection wavelength was 240 nm with a 4 nm bandwidth. Peak identities were established using commercial TPA *≥*98% purity (CAS: 100-21-0, Sigma-Aldrich), MHET *≥*95% purity (CAS: 1137-99-1, Advanced ChemBlocks), and BHET *≥*95% purity (CAS: 959-26-2; Sigma-Aldrich). Data analysis was performed in R using the chromatographR package and analyte abundance was determined by measuring absorbance peak area at 240 nm. TPA and its ester derivatives (BHET and MHET) have similar extinction coefficients at 240-244 nm [69] and purified BHET dimer (see below) consistently gave peak areas that were 3.5 times smaller than pure BHET across a range of analyzed amounts (Extended Data Fig. 3 f-g). Therefore, product formation was expressed as a ratio between the peak area of the enzymatic reaction products TPA, MHET and 3.5-times the peak area of BHET dimer, relative to the sum of all peaks (TPA, MHET, 3.5 times BHET dimer, and BHET). Finally, the relative abundance of each analyte was normalized to the cell concentration at the time of BHET addition in each reaction determined using a Beckman-Coulter Counter Z1 equipped with a 100 micron aperture tube.

For BHET conversion measurement from halo zones on YPD+BHET agar plates, colonies were washed off of the plates and white halos or clearings were excised using a pipet tip with a 1-2 mm diameter. The agar plugs were soaked in 250 ul of 100% DMSO for 24 hours at room temperature prior to centrifugation at 15,000 rpm for 3 minutes. The supernatant fraction was further diluted 5-fold prior to HPLC analysis (see above) or LC-MS (see below). Samples for LC-MS analysis were prepared in low-bind tubes and centrifuged for 20 minutes at 15,000 rpm prior to analysis to prevent any transfer of large particles into the instrument.

### 3.10 BHET dimer purification

20 *µ*moles of BHET from a 1M BHET stock solution (CAS: 959-26-2, Sigma-Aldrich) which contained BHET dimer (O,O*^′^*-(ethane-1,2-diyl) bis(oxy(2-hydroxyethyl)carbonyl)terephthalate) as impurity was fractionated by HPLC using the aforementioned fractionation protocol. Fractions between 28-29 minutes and 33.9-35 minutes were collected from the waste line of the instrument prior to being dried at 60°C for 24 hours to isolate BHET and BHET dimer, respectively. The weight of the dried fraction was then measured on a high-precision scale and the dried fraction was resuspend in 100% DMSO to a final concentration of 1 M. Fraction purity was verified by HPLC analysis across a range of concentrations (1 mM – 125 *µ*M, Extended Data Fig. 3e-f).

### 3.11 Liquid chromatography - Mass spectrometry analysis

Halo zones from under yeast colonies were extracted from agar plates as described above. An Agilent 1260 Infinity II with 6545 LC/QTOF mass spectrometer was used to analyze the samples in positive ionization mode with Dual AJS electrospray ionization (ESI) equipped with Agilent ZORBAX Eclipse Plus C18 column (2.1×50mm, 1.8-*µ*m particles) and ZORBAX Eclipse Plus C18 guard column (2.1×5mm, 1.8-*µ*m particles). LC parameters were as follows: injection volume 2*µ*L preceded with a 4*µ*L needle wash with sample, autosampler chamber temperature 20°C, column oven temperature 40°C. Mass spectrometry parameters were as follows: gas temperature 320°C, drying gas flow 10 L/min, nebulizer 35 psi, sheath gas 350°C at 11 liters per minute, VCap 3500V, Nozzle voltage 1000V, fragmentor 125V, skimmer 65V. The solvent gradient with a flow of 0.5 ml per minute started with 99% mobile phase A (Optima LC/MS H_2_O+0.1% Formic Acid, Fisher Chemical P/N LS118-4) and 1% mobile phase B (Optima LC/MS Acetonitrile+0.1% Formic Acid, Fisher Chemical P/N LS120-4), kept for five minutes, increased linearly to 100% B at 5 minutes, followed by five minutes at 100% B, then back to 1% B over 2 minutes and finally held at 1% B for an additional 5 minutes. The post-run time was two minutes (instrument conditioning at 99% mobile phase A). The raw data was analyzed using Agilent MassHunter Qualitative Analysis 12.0. Counts of molecules with mass-to-charge (m/z) ratios specific to MHET, BHET and BHET dimer were collected, and the area under the curve of each peak was calculated to determine the abundance of each molecule. MHET and BHET dimer abundance was expressed relative to spectral counts obtained for BHET in each sample. A putative BHET dimer structure was inferred from the m/z ratio.

### 3.12 Discovery and characterization of HHV-6 reactivation

For all HHV-6 analyses, we utilized Logan-Search using the AF157706 reference genome and transcriptome, reflecting the HHV-6B strain, which is endemic outside of Sub-Saharan Africa. For selecting HHV-6 transcripts to query in Logan-Search, we prioritized the HHV-6 genes annotated as ‘late’ that encode proteins essential for viral assembly and release of particles. In doing so, our search prioritized libraries containing viral expression consistent with full reactivation (rather than latency or early reactivation). The full RefSeq transcripts for U83 and U91 were input to Logan-Search and queried against the full set of human RNA-Seq datasets. Once these accessions were identified from the full search, individual SRR files were downloaded and further analyzed for modality-specific analyses. For bulk RNA-seq libraries, kallisto[70] in quant mode was used with the HHV-6 transcriptome as previously described [22]. For single-cell analyses of the organoid system, processed human counts matrices were downloaded from GEO and further annotated with the kallisto bus single-cell counts for HHV-6. For ChIP-seq libraries, raw .fastq files were re-mapped to the HHV-6 reference genome using bwa [71] with downstream analyses conducted using GenomicAlignments. Metadata, including the clinical trial identifier, input cell type, and donor identity, was pulled from the SRA metadata annotations per BioProject.

To mitigate sources of confounding for downstream analyses (i.e., errant, non-HHV-6 detection), we performed a series of stringent filters for read quantification. First, to minimize the possibility of multi-mapping transcripts, we excluded the HHV-6 DR1 transcript that possesses high homology with human transcripts [22]. Second, to mitigate the possibility of HHV-6 contamination (either environmental or index hopping), we a limit of detection per library requiring (a) a minimum of 3 unique genes and (b) minimum of 10 unique sequencing reads to call a library positive. Finally, for libraries with high viral reactivation, we verified single nucleotide diversity comparing mutations with a minimum cover of 10x per library and allele frequencies exceeding 90% in one library and less than 10% in the other Extended Data Fig. 6d). Libraries meeting these criteria were further processed to estimate the viral RNA counts per million defined as the non-DR1 HHV-6 transcripts divided by the total sequencing reads per library. Hence, we emphasize that these additional measures yield a conservative estimate of HHV-6 abundance in these libraries.

### 3.13 Gene calling and protein clustering

Protein-coding genes were predicted in all assembled Logan contigs using Prodigal [72] (version 2.6.3) in metagenomic mode (-p meta). The predicted proteins were then divided into ‘human’ vs. ‘other’ based on SRA metadata associated with their contigs through their SRA accessions and into ‘complete’ vs. ‘partial’, based on Prodigal’s output. In all, three subsets were obtained: 31.2 billion “human-complete”, 109.4 billion “other-complete”, and 304.7 billion “human-partial” protein sequences. We excluded the “other-partial” set because partial predictions are more error-prone and because this set alone contained nearly one trillion sequences that did not cluster well.

These subsets were then clustered using Linclust [73] (commit 62a2ad) on AWS Batch/EC2; the input was streamed from Amazon S3 and partitioned into fixed-size line chunks sized to instance memory (up to 1.5 billion proteins per chunk). Jobs ran as Batch array jobs on x2gd.metal nodes with a 700G split-memory limit (--split-memory-limit 700G).

Linclust was run in several cascaded rounds [74]. First, we removed near-duplicate protein fragments at 90% sequence identity (--min-seq-id 0.9) with target coverage 90% (--cov-mode 1 -c 0.9) until convergence, requiring 2, 3, and 3 rounds for the human-complete, other-complete, and human-partial sets, respectively. This step yielded 0.154B, 7.4B, and 10.8B clusters, each of which with its representative sequence, termed Logan90.

Next, the Logan90 databases were further clustered at 50% identity with the same coverage criterion until convergence, requiring 1, 3, and 2 rounds for the human-complete, other-complete, and human-partial sets, respectively. To ensure that dividing sequences into chunks during the final 50%-identity clustering rounds (other-complete rounds 2–3; human-partial rounds 1–2) did not prevent the detection of sequences belonging to the same cluster, we shuffled sequences between chunks. The final representative sets, termed Logan50 databases, contained 0.07B, 3B, and 1.8B sequences, corresponding to overall reductions of 99.8%, 97.3%, and 99.4%. Altogether, we reduced 445.3B initial sequences to 4.87B representative sequences, a total reduction of 98.9%.

#### 3.13.1 Logan50 clustering enhances protein structure prediction by increasing multiple sequence alignments (MSAs) diversity

To demonstrate the utility of Logan50 for downstream applications, we tested whether they can improve protein structure predictions with AlphaFold2/ColabFold [33], focusing on viral proteins whose structures are hard to predict with the default ColabFold DB. The Big Fantastic Virus Database [75] (BFVD) comprises over 351,000 viral protein structures, generated by augmenting ColabFold default MSAs with homologous proteins identified in Logan’s contigs. In the BFVD paper, identification of homologs was carried out over the entire 0.9 petabases of Logan contigs. Here, we tested whether the reduced Logan50 dataset could achieve comparable performance while reducing the search space by approximately 3,500-fold compared to using the entire Logan corpus.

We obtained 100 viral proteins from BFVD, previously characterized by poor-quality MSAs using the default ColabFold DB. For each protein, we generated MSAs using MMseqs2 (mmseqs search -a -s 8.5 --num-iterations 2 --max-seqs 1000) against the other-complete Logan50 database, and compared them to the default ColabFold MSAs in terms of MSA quality, measured by the number of effective sequences (Neff) with hhmake (Fig. 5b, left) and structural prediction (Fig. 5b, right) quality, measured by the pLDDT metric. Protein structure prediction using either default MSAs or Logan50 MSAs was done by AlphaFold2/ColabFold (colabfold batch --num-models 1 --model-order 3).

We observed substantial improvements in both metrics. Average Neff increased from 2.19 (baseline) to 4.89 (Logan50 MSAs), while mean pLDDT scores improved from 46.7 (“very low”) to 88.6 (“high”). Combining ColabFold and Logan50 MSAs further increased these values to a Neff of 5.18 and mean pLDDT of 89.02, approaching the improvement achieved with the full Logan corpus plus ColabFold sequences in the original BFVD paper (mean Neff = 4.39, mean pLDDT = 92.88). Thus, on par significant gains in structural prediction can be achieved efficiently using the thousand times smaller Logan50, with substantial reductions in computational requirements.

### 3.14 Expanding Obelisk Diversity with Logan

#### 3.14.1 Reconstructing the Original Obelisk Database

A total of 1,744 Obelisk clusters (80% nucleotide clustering threshold), representing 7,202 circular genomes, were taken from the initial Obelisk study [9]. These 1,744 centroid were screened for false positives using several sequence alignment, structural homology and HMM verification steps (Data Availability, Obelisk Data Methods). The final clustering analysis resulted in 1,284 backward-compatible 60% Oblin-1 clusters and 1,965 90% clusters. Each of the 1,284 60% centroids also functioned as a 90% centroid, maintaining consistency in the database. This re-annotated database is designated as ‘Obelisk DB Legacy’.

#### 3.14.2 First Obelisk Expansion

The Obelisk DB Legacy 1,744 Oblin-1 centroid sequences were used as query sequences for alignment to all Logan contigs using minimap2 (Section 3.2). The results of the Logan search were retrieved and split into circular and non-circular sequences by screening for 30-mer repeats at the ends of the Logan contigs. For circular sequences, the 30-mer repeat was trimmed, yielding 12,690 circular and 312,728 non-circular contigs.

Concurrently, the 7,195 Oblin-1 sequences from ‘Obelisk DB Legacy’ were clustered at 95% identity using USEARCH cluster fast, resulting in 2,170 clusters. Using Clustal Omega [76] and HMMER [63], a Hidden Markov Model was constructed from this clustered dataset. The circular sequences from the Logan output underwent ORF calling using EMBOSS getorf with circular genome parameters (getorf-circular Yes), and the resulting ORFs were incorporated into the ‘Obelisk DB Legacy’ HMM model through an iterative search and alignment protocol (Data Availability, Obelisk Data Methods).

This procedure resulted in a comprehensive Obelisk database containing 117,838 Oblin-1 proteins, along with a high-quality MSA comprised of 45,170 Oblin-1 sequence alignments. All 117,838 Oblin-1 proteins were verified to contain the Domain A motif, and were aligned against the BLAST nr database (May 2024) using DIAMOND BLASTp [11] with parameters –masking 0 –unal 1 –sensitive -c1 -k1 -b 5 –threads 16. This analysis confirmed that no Obelisk sequences produced significant hits in the BLAST nr database, indicating that all sequences represent completely novel genetic elements.

#### 3.14.3 Construction of Obelisk DB v1

The original Obelisk sequences from ‘Obelisk DB Legacy’ were excluded from the 117,838 Obelisk sequences, yielding 110,643 novel sequences. These novel sequences were aligned to the 1,284 60% centroid sequences from the Legacy Database using USEARCH usearch global with 60% identity threshold (usearch -usearch global -id 0.6). Sequences that aligned within known 60% clusters were subsequently aligned to the corresponding 90% centroid sequences within their respective 60% clusters using USEARCH usearch global with 90% identity threshold (usearch -usearch global -id 0.9).

This two-step alignment procedure classified all sequences into three distinct categories: (1) members of known 60% and 90% clusters, (2) members of known 60% clusters that represent novel 90% clusters, or (3) members of novel 60% clusters. Using the alignment identities and closest matches identified by USEARCH, sequences were assigned to their appropriate clusters.

For novel 60% or 90% clusters, centroid selection prioritized circular sequences, followed by sequence length as the secondary criterion. In clusters lacking circular sequences, the longest sequence was designated as the representative centroid. Metadata, including amino acid sequence, nucleotide sequence, circularity status, SRA accession numbers, BioProject information, and additional relevant data, was appended to each database entry. The resulting database was designated ‘Obelisk DB Logan v1’.

#### 3.14.4 Database Refinement with Logan v1.1 Contigs

As Logan contigs were updated from v1 to v1.1, we also updated the Obelisk database. Centroids from the 90% Obelisk clusters were extracted and aligned to Logan v1.1 contigs using minimap2 (Section 3.2). The results were processed using the same methodology described in the previous section. Circular and non-circular sequences were separated, and *k*-mer repeats were removed following the established protocol. The v1.1 Obelisk database was constructed as follows. Beginning with the original 7,195 sequences and the previously established high-quality 45,000-sequence MSA, the same iterative alignment methodology with tapered e-value cutoffs was executed. Circular sequences were processed first, followed by non-circular sequences. In total 67,454 Oblin-1 sequences were captured.

BLAST alignment analysis revealed no significant hits for all proteins except one sequence, which showed a very distant, low e-value alignment. The constructed HMM model was used to query all 67,454 sequences, and only 550 Oblin-1 sequences exhibited alignment e-values greater than e-5 to the model; these sequences were retained to preserve diversity. This database was designated ‘Obelisk DB Logan v1.1’.

Comparative analysis using DIAMOND BLASTp between versions 1 and 1.1 confirmed that all sequences present in version 1 were accounted for in version 1.1, despite the total number of Obelisk sequences being approximately halved. This reduction can be attributed to the differences in average sequence lengths between the two databases. Version 1 had an average amino acid length of 136.7 residues, while version 1.1 had an average length of 186 residues. This increase in sequence length explains the decrease in individual sequence counts, while maintaining comprehensive coverage.

The same methodology described in ‘Construction of Obelisk DB v1’ was applied and the complete ‘Obelisk DB v1.1’ was assembled with all associated metadata incorporated.

### 3.15 Genomic Expansion of P4 Phage Satellites

We first gathered all RefSeq protein sequences for each of the seven core components in a P4 phage satellite [34] (Counts: alpha: 1911, ash: 1995, alpA: 2008, Sid: 2041, Delta: 2089, Psu: 1891, Integrase: 2097). Sequences were clustered for each core component individually via UCLUST v11.0.667 cluster fast [60] with identity threshold set to 0.9, then cluster representatives from each of the core components were aligned against Logan v1.1 contigs via DIAMOND BLASTX, as per Section 3.2, with sensitive mode and e-value minimum of 1e-8, for a total of 107.6 million putative P4 satellite contigs. Contigs were next refined with HMMER v3.4 [63], by 6-frame translating each of the putative P4 contigs with seqkit v0.10.0 translate [49], then running HMMER hmmsearch on each of the translated sequence against each of the core component protein databases. A total of 210,895 contigs contained a hmmsearch hit for each core component with a minimum e-value of 1e-5 and a coverage of at least 40% of the database sequence. Finally, these contigs were clustered with MMseq2 v15 [77] with minimum identity threshold of 0.999 and coverage of 0.8, curating 16,215 cluster representatives. Proteins were detected within each representative with Prodigal [72] (version 2.6.3) and P4 phage satellites were detected and characterized with SatelliteFinder v1 [34] with default parameters.

For the pangenome curve, proteins from all RefSeq P4 phage satellites of types A, B, or C were clustered with MMseqs2. Clustering was then repeated with each cluster representative and proteins from Type A and B Logan contig containing less than 30 genes (15,653 contigs). Any cluster containing only Logan contigs was considered a new gene family. For the whole-genome reciprocal relatedness (wGRR) analysis, full proteomes from the 16,653 Logan P4 contigs and the 3,437 RefSeq P4 genomes were extracted by detecting the first and last protein in the P4 satellite region. A wGRR matrix was formed by calculating the fraction of bi-directional best hits weighted by the sequence identity for each genome pair in an all- vs-all fashion. Hierarchical clustering was performed on the matrix and the corresponding heatmap was rendered with with seaborn’s clustermap function, using the Ward clustering algorithm. All scripts can be accessed at https://github.com/kdcurry/P4-logan.

### 3.16 Plasmid identification and clustering

We identified 3,885,511 circular contigs (*≥* 1% kb) from version 1.0 assembly graphs of 252,507 samples, including bacterial, archaeal, and metagenomic datasets (code available at https://gitlab.pasteur.fr/rchikhi_pasteur/logan-circles). Contigs were processed with tr-trimmer (version 0.1.0, parameters: -c -x -l 31) to discard sequences with low-complexity repeats spanning *>* 50% of terminal 31-bp repeats and to trim these repeats from the 3’ ends. To mitigate gene truncation, sequence breakpoints were shifted to intergenic regions (code available at https://github.com/apcamargo/reorient-circular-seq). Plasmids were then identified using geNomad [78] (version 1.8.1, database version 1.7, end-to-end command, parameters: --enable-score-calibration --max-fdr 0.01). To minimize false positives, we only kept plasmids that met two criteria: (1) encode at least one protein matching a HMM from a curated set of 193 plasmid hallmark protein profiles (see Data Availability); (2) encode no more than two proteins matching HMMs of near-universal single-copy orthologs from the BUSCO v5 [79] odb10 [80] datasets of *Bacteria* and *Archaea*. Gene prediction was carried out using pyrodigal-gv [81, 78] (version 0.3.2), and protein sequence matching to HMMs was performed using PyHMMER’s [82] (version 0.10.15) hmmsearch function, applying gathering cutoffs to the HMMs of plasmid hallmarks (parameter: bit_cutoffs=“gathering”) and BUSCO bitscore cutoffs to the HMMs of near-universal single-copy orthologs. Identified plasmids were assigned to replicon families using MOB-typer [83] (version 3.1.9). AMR genes were annotated using AMRFinderPlus [84] (version 4.0.3), and AMPs were identified with Macrel [85] (version 1.5.0).

To cluster plasmids, we first computed pairwise sequence similarities using BLAST [57] (version 2.16.0, parameters: -task megablast -evalue 1e-5) and the anicalc.py script from CheckV [86] to compute pairwise similarity metrics. We then constructed a similarity graph connecting pairs of plasmids exhibiting sequence identity *≥* 90% and bidirectional alignment coverage *≥* 90%, and clustered the plasmids using the pyLeiden [87] tool.

We evaluated Faith’s phylogenetic diversity of selected replicases and relaxases (Data Availability, Plasmid PD Table) from newly identified plasmids and complete plasmids retrieved from PLSDB (release 2024 05 31 v2) and IMG/PR. Genes encoding these proteins were identified using hmmsearch, and multiple sequence alignments were generated with PyHMMER’s hmmalign function (parameters: trim=True, all consensus cols=False). These alignments were then trimmed with the gappyout algorithm from PytrimAl [88] (version 0.8.0) and phylogenetic trees were inferred with FastTree [89] (version 2.1.11).

We surveyed plasmid presence across all samples by mapping contigs from version 1.1 assemblies to plasmid cluster representatives using minimap2 (version 2.28, parameters: -x sr --sam-hit-only -a). A plasmid cluster was considered present in a sample if *≥* 75% of its length was covered by alignments from that sample.

### 3.17 Antimicrobial resistance genes discovery and analyses

We analysed the presence of antimicrobial resistance (AMR) genes across 26.7 million SRA accessions via the Logan v1.1 contigs. AMR gene hits were identified by aligning the CARD nucleotide database (version 3.3.0) to Logan contigs using minimap2, as per Section 3.2 (see Data Availability). Alignments were filtered to contain only those sequences with *>*100 bp length and *>*80% identity. SRA metadata and extended geolocation data (see Data Availability) were used to classify information on CARD alignment SRA accession hits. For analyses based on organism classification, datasets were classified as organism type Metagenome if SRA metadata field organism contains the string “metagenome”, or if the field librarysource contains “METAGENOMIC” or “METATRANSCRIPTOMIC”; otherwise, accessions were classified as Isolate. Metagenome categories were classified according to the organism and librarysource fields, dividing it into the 6 top categories: human, livestock, soil, marine, freshwater, wastewater. Plasmid metadata was extended analogously to SRA samples. For Extended Data Fig. 7d-g), the SRA-CARD alignment dataset was filtered to include only samples with known collection dates, geolocation, metagenomic origin, and those more likely to contain whole genome/transcriptome assay types (WGS, WGA, RNA-Seq, etc.). All code and datasets can be accessed in: https://github.com/mmontonerin/logan_AMR

### 3.18 Comparison of Logan metagenomes with GenBank WGS via FracMinHash sketches

FracMinHash sketches were created with sourmash [90] for each Logan unitigs accession whose SRA metadata information indicated it was a metagenome (total of 4,792,069 accessions). We used a *k*-mer size of 31 and a scale factor of 1,000. The resulting sketches, comprised of 449,087,511,713 hashes with duplicates, were then placed in a duckDB database [91]. Code to process these data are available at: https://github.com/KoslickiLab/ingest_logan_yacht_data.

We then crawled the GenBank Whole Genome Shotgun (WGS) FTP server, downloaded and sketched each *.fsa nt.gz assembly with sourmash using the same *k*-mer size of 31 and scale factor of 1,000. Of the 2,055,047 *.fsa nt.gz files discovered on this FTP server, 5 led to HTTP error code 404 when attempting to download them. Signatures were stored as compressed *.sig.zip archives. We extracted all 64-bit FracMinHash hash values, a total of 32,701,966,322 hashes with duplicates, and partitioned them by a low-bits bucket function to enable external-memory de-duplication. We then reduced each bucket to its exact set of 8,679,649,739 unique hashes and wrote a partitioned Parquet dataset which we ingest into DuckDB. To compare against the Logan metagenome sketches, we attached the Logan metagenome FracMinHash sketches, and again bucketed into Parquet files and computed set differences with bucket-wise anti-joins in DuckDB to report the number of 31-mers (appearing twice or more) unique to each dataset: 32,287,730,882 in the Logan metagenomes not in GenBank WGS, and 7,020,467,844 in GenBank WGS not in the Logan metagenome sketches. Since a scale factor of 1,000 was used to form the sketches, multiplying these number of hashes by 1,000 results in the estimated total number of 31-mers, each appearing twice or more in each dataset. All code can be accessed at: https://github.com/KoslickiLab/GenBank_WGS_analysis.

### 3.19 Geographic Metadata Extraction, Inference and Enrichment

99.7% of SRA submissions are associated with a BioSample https://www.ncbi.nlm.nih.gov/biosample record. BioSamples contain zero or many attribute [name, value] tuples with submitter-supplied metadata that describes the biological sample. A full XML dump of the BioSample database was retrieved on June 28, 2024, comprising 39,448,576 records with 568,885,433 attribute entries. A subset of attribute names likely to contain geographical information was extracted using a large language model (gpt-3.5-turbo-0125) combined with manual curation of the most frequently occurring ones. The distribution of attribute names is heavily skewed and long-tailed: the eight most common names account for 92.84% of all entries identified as containing geographical information.

Attribute values corresponding to this subset of attribute names were then used to infer geographic coordinates. A deep learning classifier partitioned the values into three distinct categories: values likely to contain numerical latitude and longitude pairs (coordinates); values likely to reference a location by its common name (place names); and values which were explicitly annotated but whose meaning is uninformative, e.g. “N/A”, “null”, “undefined”, etc. Coordinates were resolved directly to points on Earth under the WGS 84 coordinate system. Place names were converted to coordinates using three different geolocation services (Azure Maps https://azure.microsoft.com/en-us/products/azure-maps; and Esri, HERE through AWS Location https://aws.amazon.com/location/). To quantify confidence in these derived coordinates, a score of 0 to 6 was assigned depending on whether the locations returned by the three services were within 8 km of each other (up to 3 points), and whether they were found within the political boundaries of the same country (up to 3 points). Values classified as uninformative were discarded. In total, geographic coordinates were obtained for 26,962,465 (68.34%) of BioSample entries.

The geographic dataset was further enriched by cross-referencing it with the following publicly available resources: *ASTER Global Digital Elevation Model* https://cmr.earthdata.nasa.gov/search/concepts/C1711961296-LPCLOUD.html to extract elevation in meters above mean sea level; *World Administrative Boundaries* https://public.opendatasoft.com/explore/dataset/world-administrative-boundaries/export/ to assign political boundaries such as countries and regions; and *WWF Terrestrial Ecoregions of the World* https://www.worldwildlife.org/publications/terrestrial-ecoregions-of-the-world to classify BioSamples into 14 distinct biomes characterized by their unique biodiversity and environmental conditions.

## 4 Data Availability

Logan is publicly available [37] on AWS Registry of Open Data at https://registry.opendata.aws/pasteur-logan/. There are no egress charges and anonymous access is permitted. A data access tutorial is provided at https://github.com/IndexThePlanet/Logan. *PETadex* data can be found at https://github.com/ababaian/petadex. AMR datasets can be found at s3://logan-pub/paper/AMR. Obelisk data and code can be found at s3://logan-pub/paper/Obelisk, and the Obelisk Data Methods is methods.pdf within this folder. Plasmid sequence and metadata can be found at s3://logan-pub/paper/plasmids, and the Plasmid PD Table is plasmid_phylogenetic_diversity.xlsx within this folder.

## 5 Acknowledgments

We are grateful to the entire team managing the NCBI SRA / EBI ENA and the biology community for data sharing. We thank Dorian Schaal, Adrien Laińe, Coral Kennett, Candi Jeronimo, Dave Maurer, and Morgan Lim from Amazon Web Services (AWS) for support. Thomas Menard, Stéphane Fournier and the HPC Core Facility from Institut Pasteur for IT support. Peter Schmiedeskamp, Chris Stoner, Erin Chu and Beryl Rabindran from AWS Registry of Open Data for data hosting. Ryan Connor and Yuriy Skripchenko from NCBI for assistance with SRA operations. Karen McGregor, Jerry Morey and Venkat Malladi from Microsoft for Azure support. Matthieu Falce for custom AWS tooling. Stephen Nayfach for feedback on the analysis of plasmid genomes. Karin Steffen and Zamin Iqbal for feedback on the manuscript. Administrative support was provided by Mélanie Ridel, Löıc Orellou, and Florence Percie du Sert.

## 7 Contributions

R.C. and A.B. conceived and led the study. R.C., G.A., M.H., A.K. and B.R. designed and implemented the Logan assemblage cloud infrastructure. A.K. developed the f2sz software. R.C. designed and implemented the Logan contigs mining analyses. T.L. and P.P developed Logan-Search. A.B., R.L-K., R.C.E, J.S., R.C., K.S., P.J.R, and G.W.B analyzed the plastic-active enzymes. C.A.L. analyzed the viral reactivation. M.S. and J.L. clustered the Logan proteins, and M.S. analyzed the viral proteins case study. R.F. created protein embeddings. P.G. analyzed the Obelisks. K.D.C., J.A.M.S and E.P.C.R. analyzed the P4 satellites. A.P.C., C.J.M. and M.B.F. analyzed the plasmids. M.M-N., D.P.A. and S.M. designed the AMR study, M.M-N. analyzed the AMR in Logan contigs, M.M-N. and A.P.C. analyzed the AMR in plasmids. D.K. analyzed Logan metagenome contigs and WGS sketches. A.M-T. created the SRA geographic metadata dataset. R.C., A.B., R.C.E., C.A.L, P.P., M.S., R.L-K., A.P.C., M.M-N., P.G., K.D.C., E.P.C, D.K., P.M., and A.M-T. wrote the manuscript.

All authors contributed to, and approved the manuscript.

## 9 Materials & Correspondence

Correspondence and requests for materials should be addressed to Rayan Chikhi or Artem Babaian.

## Extended Data

**Extended Data Fig. 1:**
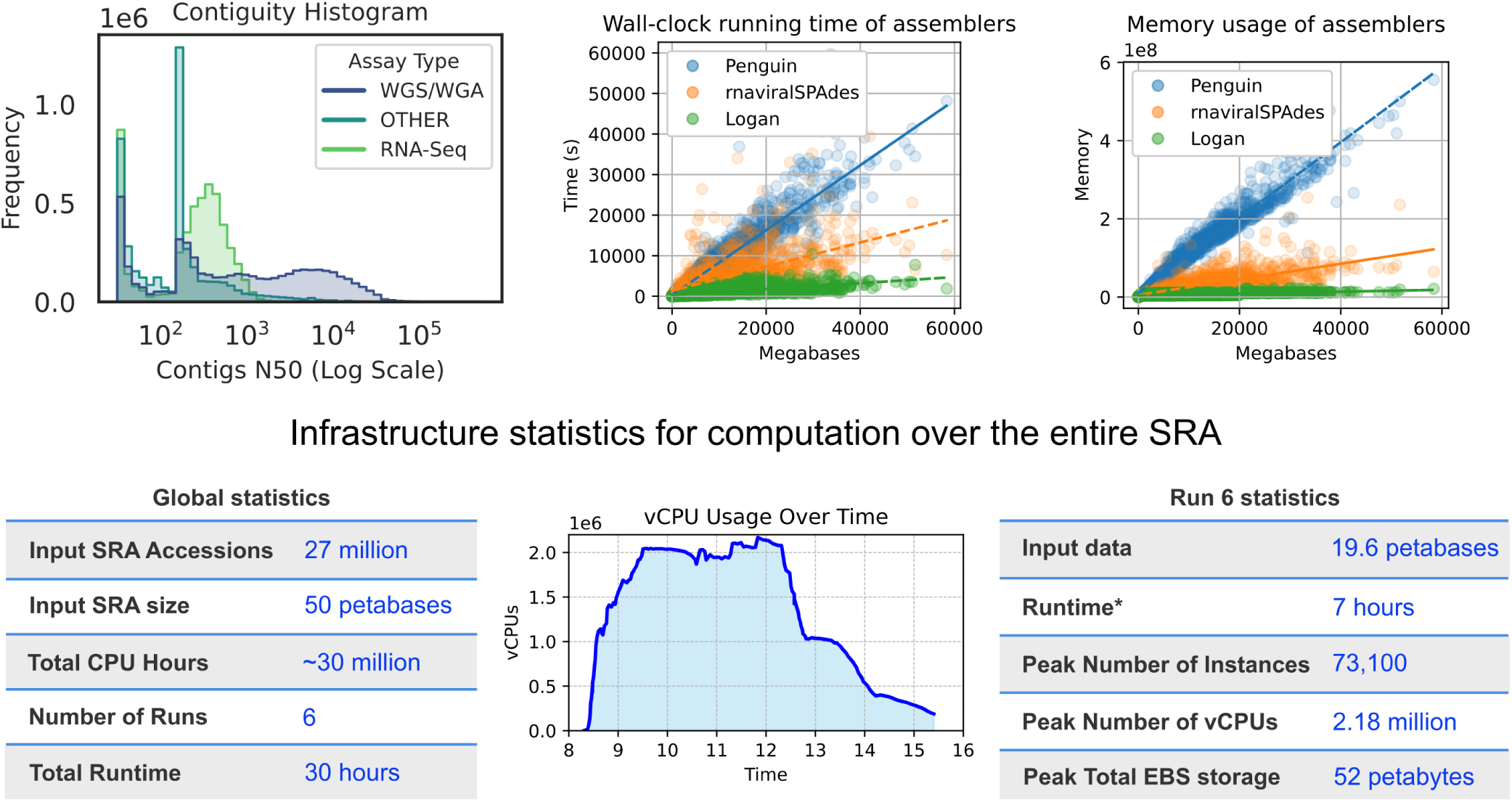
Logan assembly performance and computational statistics for processing the entire SRA. This figure details the performance benchmarks of the Logan pipeline and quantifies the cloud computing resources used to assemble 27 million SRA datasets. (Top Left) A histogram showing the distribution of assembly contiguity, measured by contig N50, across all Logan assemblies. Assemblies are categorized by input SRA assay type, showing that Whole Genome Shotgun (WGS/WGA) samples generally produce more contiguous assemblies than RNA-Seq or other samples, as expected. (Top Middle and Right) Performance benchmarks comparing the Logan assembly pipeline to other state-of-the-art short read metagenome assembly tools (Penguin, maviralSPAdes). Logan pipeline demonstrates significantly lower wall-clock running time (middle) and memory usage (right) across a range of input data sizes, highlighting its efficiency. (Bottom) Statistics from the full-scale production run. Global statistics summarize the total compute effort, including processing 50 petabases of input data over 30 million CPU hours. The vCPU Usage Over Time plot for the main production run illustrates the dynamic allocation of cloud processors, peaking at over 2.18 million vCPUs. Run 6 statistics detail the single largest run, where 19.6 petabases of data were assembled in just 7 hours.

**Extended Data Fig. 2:**
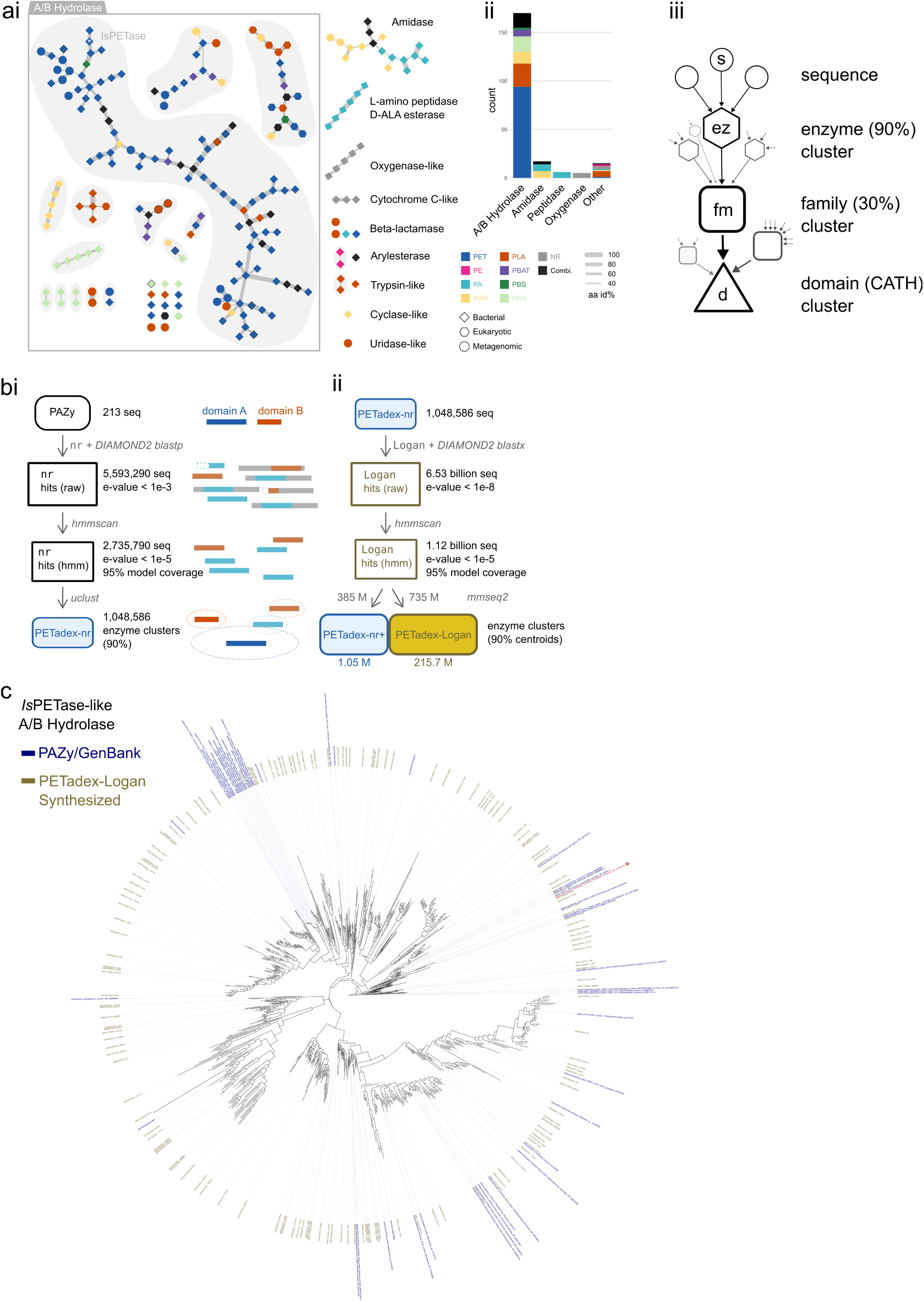
PETadex-Logan Workflow. **(a)** Characterization of the initial 213 plastic-active enzymes from the PAZy database. **(i)** A network graph showing sequence similarity between the known enzymes, colored by protein family. **(ii)** Bar chart showing the distribution of these enzymes across protein families and the types of plastics they degrade. **(iii)** Schematic of the hierarchical clustering strategy used to group sequences at the enzyme (90% identity), family (30% identity), and domain (CATH) levels. **(b)** The two-stage deep homology search pipeline. **(i)** The first stage queried PAZy sequences against the NCBI nr database. After filtering for domain integrity and clustering, this step yielded 1.05 million enzyme clusters, creating the PETadex-nr dataset. **(ii)** In the second stage, PETadex-nr was queried against the entire Logan assembled contigs, identifying 735 million novel sequences and massively expanding the diversity into the final PETadex-Logan dataset. **(c)** A phylogenetic tree of the *Is*PETaselike A/B Hydrolase clade. The tree visually demonstrates the expansion of sequence diversity uncovered by the Logan search compared to the previously known diversity from public databases like PAZy and GenBank (blue labels). Sequences selected for experimental evaluation are labeled.

**Extended Data Fig. 3:**
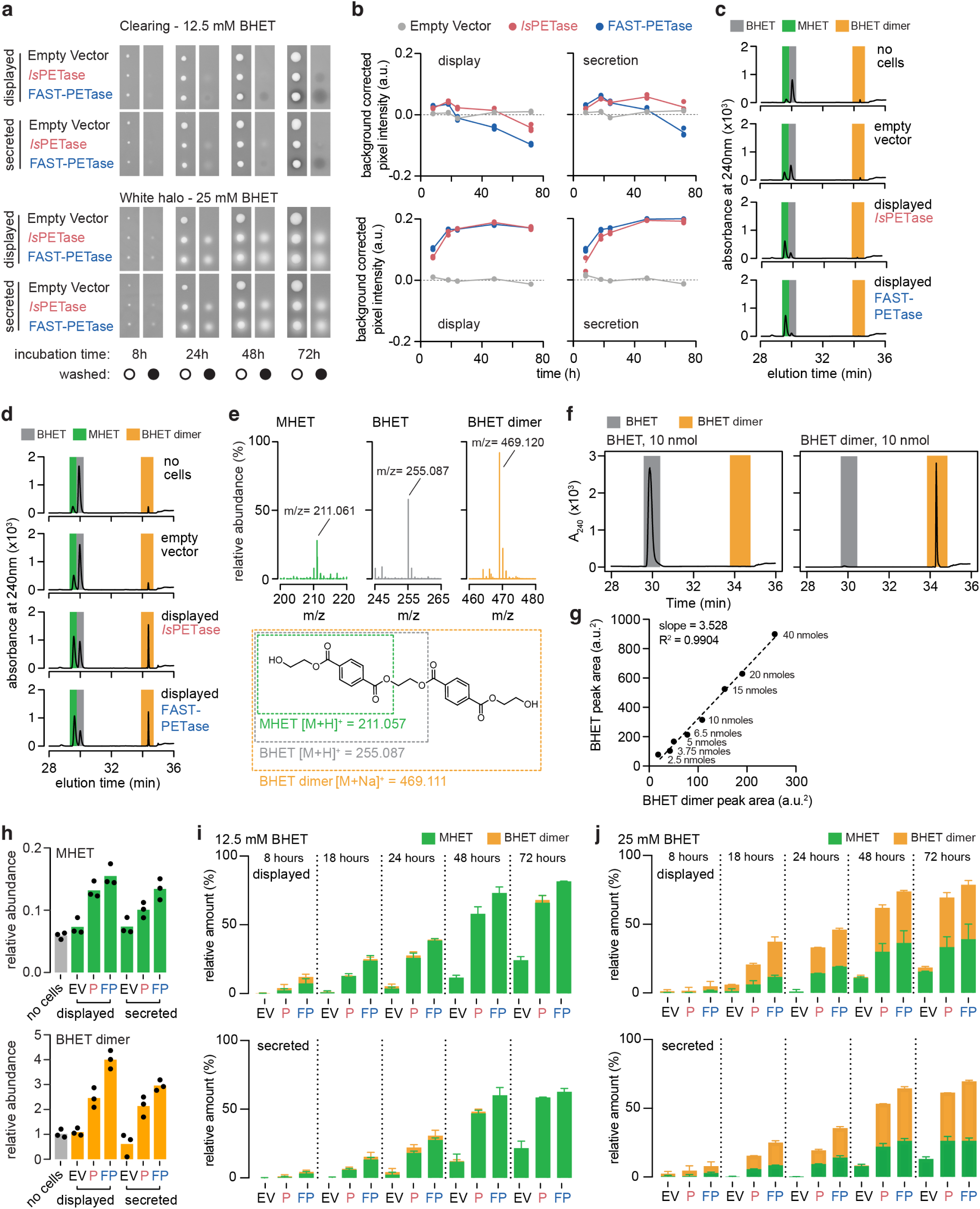
PETase Halo Assay. **(a)** Clearing and white halo detection with yeast colonies expressing surface displayed or secreted PETase enzymes. Yeast strains were robotically pinned onto YPD medium containing 12.5 mM or 25 mM BHET and incubated for 8 to 72 hours at 30 degrees Celsius. Plates were imaged before (open circle) and after (closed circle) washing colonies off the plate. **(b)** Clearing and white halo quantification from washed YPD plates containing 12.5 mM or 25 mM BHET. Pixel intensity of the colony area was measured using a custom R pipeline (see Methods) on images from (a). Data is depicted as the background normalized median pixel intensity under each colony over time for the indicated PETases; n=3. **(c)** High-Performance Liquid Chromatography (HPLC) analysis of the clearing zone identifies the MHET reaction product. Agar plugs were excised from the plates in (a) for the displayed PETases, empty vector control, and regions with no yeast cells, after 72 hours of yeast growth on 12.5 mM BHET, and dissolved in DMSO prior to HPLC analysis. Representative chromatograms are shown; *n ≥* 2. Coloured shading indicates the identity of each peak. **(d)** HPLC analysis of the white halo identifies MHET and BHET dimer reaction products. Agar plugs were excised from the plates in (a) for the displayed PETases, empty vector control, and regions with no yeast cells, after 72 hours of yeast growth on 25 mM BHET, and dissolved in DMSO prior to HPLC analysis. Representative chromatograms are shown; *n ≥* 2. Coloured shading indicates the identity of each peak. **(e)** Mass spectrometric (MS) analysis of MHET, BHET, and BHET dimer purified from a white halo extracted under a yeast colony expressing surface-displayed *Is*PETase after 24 hours on YPD plus 25 mM BHET. Representative spectra are shown with the mass to charge ratio of the most abundant component indicated; n = 3. The chemical structures of BHET, MHET and putative BHET dimer are shown along with their predicted ionized mass. **(f)** HPLC analysis of 10 nmol of HPLC-purified BHET, and 10 nmol of HPLC-purified BHET dimer. Coloured shading indicates the identity of each peak. **(g)** Correlation plot of absorbance peak areas from HPLC analysis of the indicated amounts of purified BHET and BHET dimer. The linear regression line is plotted. **(h)** MS quantification of MHET and BHET dimer from agar plug extraction. Agar plugs were obtained from an area of YPD plus BHET 25 mM with no yeast colony (no cells) or under the yeast colonies (after wash) containing the indicated constructs after 24 hours of incubation and processed as in (d). Relative abundance is plotted, expressed as a ratio between spectral counts for MHET (top) or BHET dimer (bottom) relative to the spectral counts obtained for BHET. EV: empty vector; P: IsPETase; FP: FAST-PETase. **(i,j)** Quantification of BHET conversion in clearing zones and white halos over time. Halos from the indicated strains, timepoints and BHET concentrations were processed as described in (c,d) and analyzed by HPLC. Peak area for each analyte (BHET, MHET, BHET dimer) was measured and expressed as a percentage relative to the sum of the peak areas for BHET+MHET+BHET dimer. EV: empty vector; P: IsPETase; FP: FAST-PETase.

**Extended Data Fig. 4:**
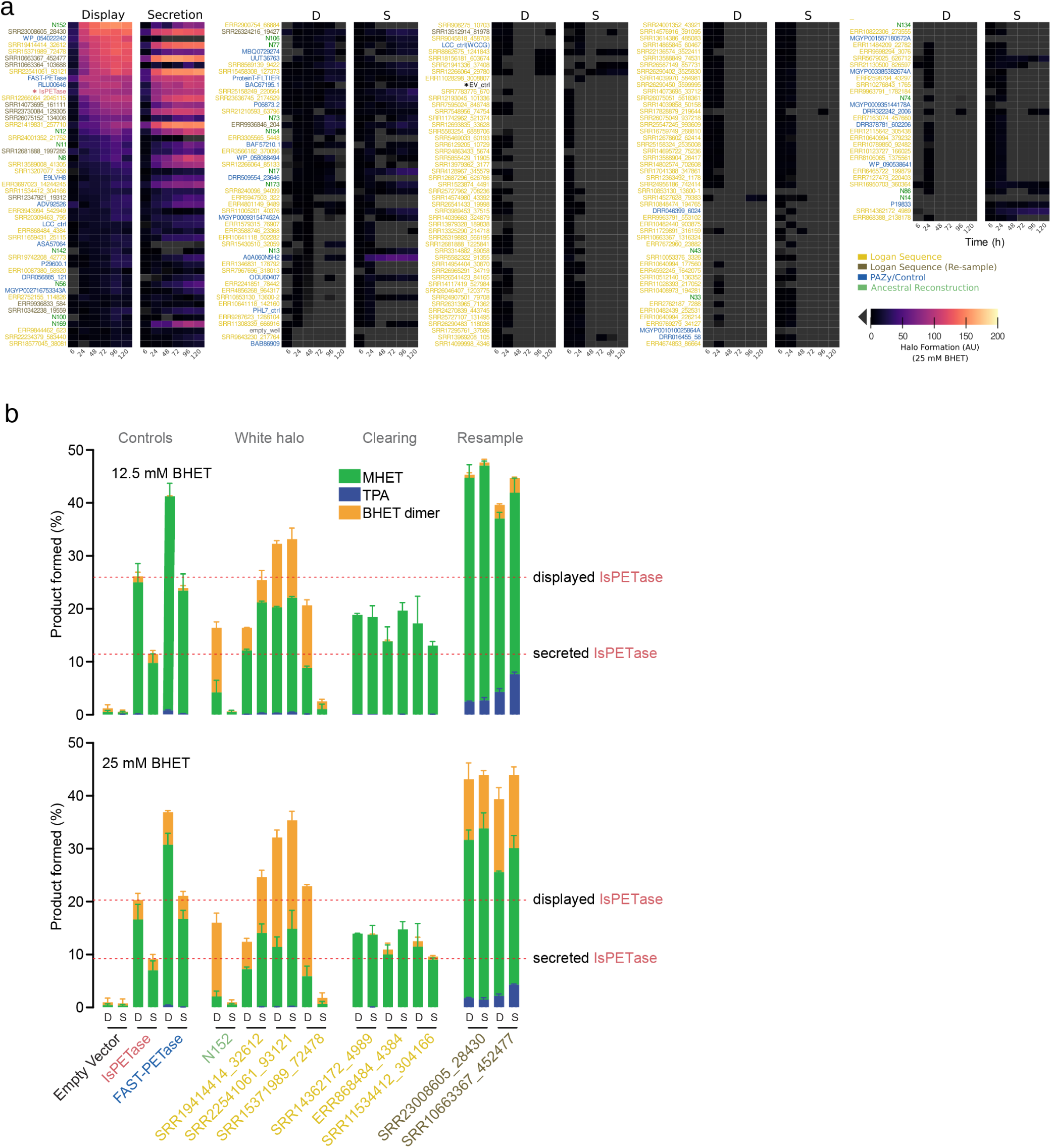
High-throughput screening and HPLC validation of PETadex-Logan enzymes. **(a)** Heatmap of the high-throughput activity screening results for Logan PETases and controls. Enzyme activity was measured as the background normalized median pixel intensity under each colony on YPD plates with 25mM BHET, at the indicated times, and in either surface displayed (D) or secreted (S) constructs. The heatmap shows the average of quadruplicate pixel intensity measurements (in arbitrary units, AU) after subtracting Empty Vector background values and scaling to approximately 100 units for *Is*PETase at 48 hours. This screen was used to identify the active candidates for quantitative analysis. **(b)** Quantification of BHET conversion in yeast strains expressing the top candidate PETase enzymes. Strains were grown to saturation in YPD medium prior to adding BHET at the indicated concentrations. BHET conversion reactions were allowed to proceed for 17 hours at 30°C, and culture supernatants were analyzed by HPLC. The peak area for each analyte (BHET, MHET, BHET dimer) was measured and expressed as a percentage of the sum of all peak areas normalized to 10^8^ cells/ml, based on the cell concentration at the time of BHET addition. D: surface displayed enzyme; S: secreted enzyme.

**Extended Data Fig. 5:**
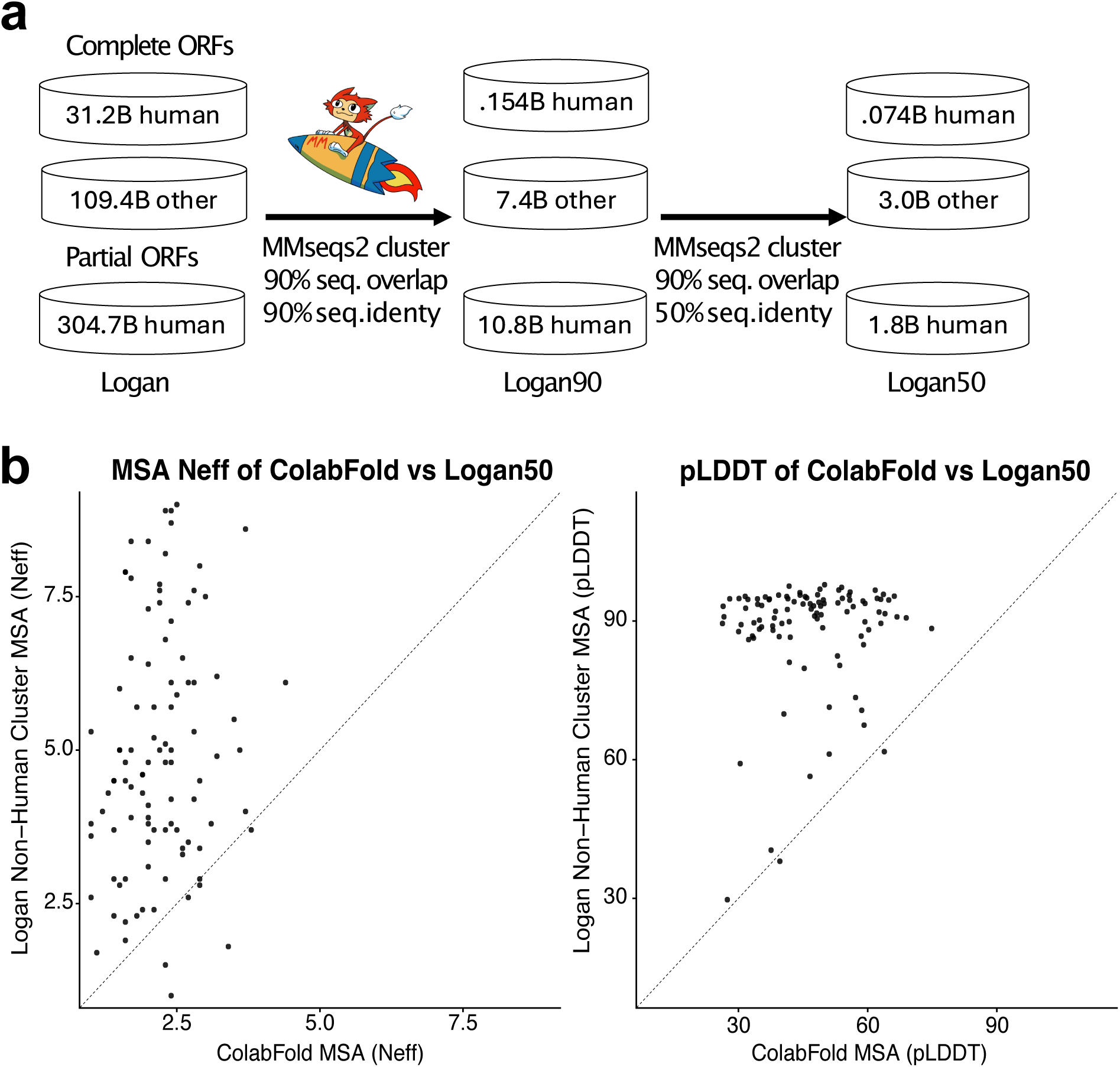
Protein clustering workflow and its application in improving multiple sequence alignment (MSA) diversity. **(a)** The workflow for creating the Logan90 and Logan50 clustered protein databases. Prodigal-predicted protein coding regions from all Logan contigs were first separated into ‘human’ and ‘other’ categories, based on SRA metadata associated with their contig and into ‘complete’ and ‘partial’, based on Prodigal’s output. The proteins were then clustered using Linclust at 90% and subsequently 50% sequence identity to create representative protein sets for sensitive homology searches. The numbers indicate billions (B) of proteins at each stage of the workflow. **(b)** A case study demonstrating the value of Logan50 for enhancing MSA diversity (Neff, left panel) and improving structure prediction quality (pLDDT, right panel) of 100 viral proteins with low-quality MSAs from the default ColabFold database. We performed sensitive, iterative profile searches against the other-complete Logan50 database (y-axis) using MMseqs2 and compared the results to those from the default ColabFold database (x-axis). In both panels, nearly all points lie above the diagonal, indicating that Logan50 yields more diverse alignments and substantially improved structural predictions.

**Extended Data Fig. 6:**
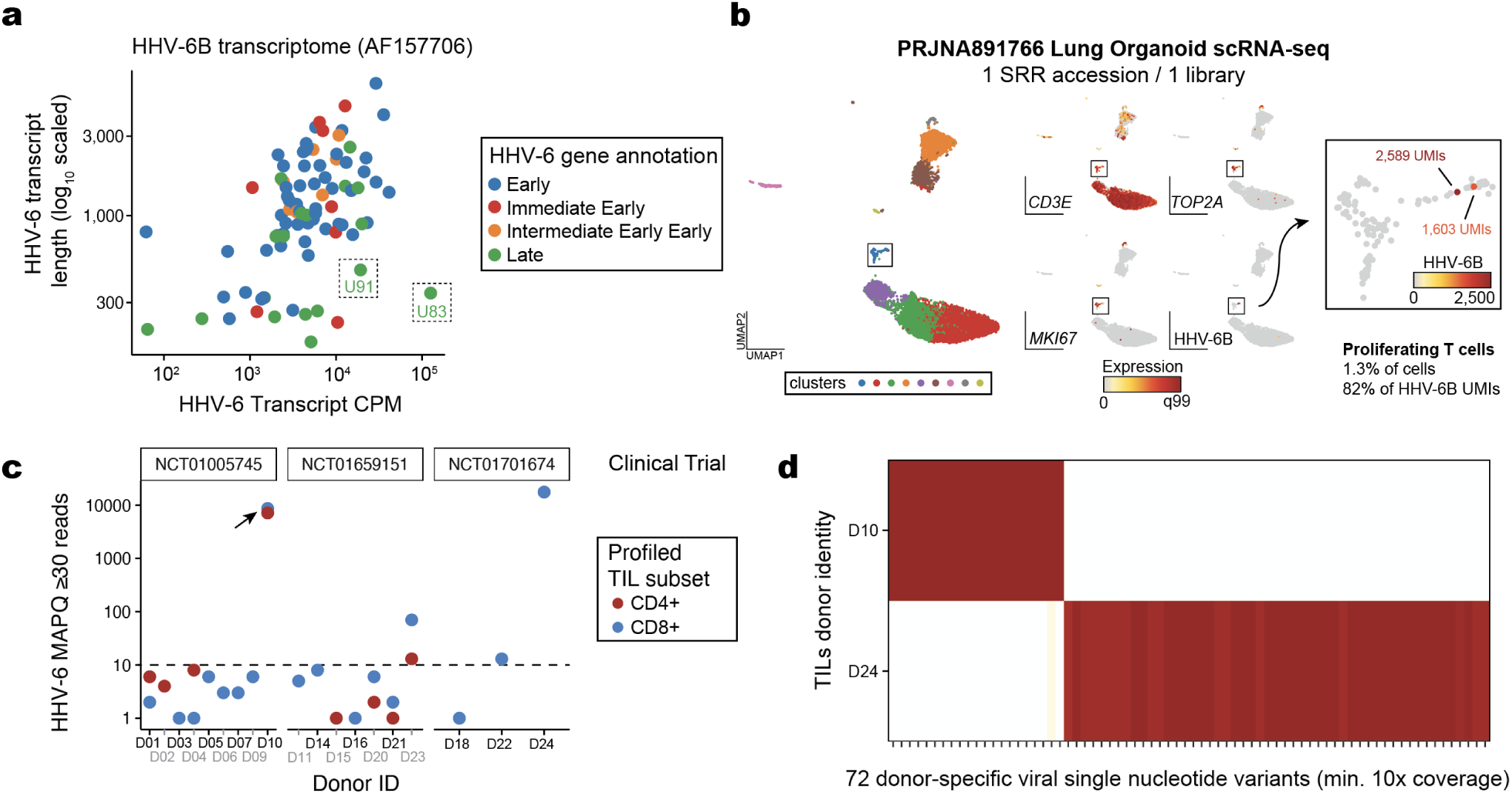
Supporting information for identification and reactivation of HHV-6 in large-scale RNA-seq datasets. **(a)** Summary of HHV-6B reference transcriptome. Viral transcript abundance was computed from a prior characterization of HHV-6 reactivation in CAR T cells (Sample 34; Day 19) [22]. Boxed genes represent selected sequences used as queries for Logan-Search. **(b)** UMAP representation of cells profiled via scRNA-seq from lung organoid culture (PRJNA891766). Panels reflect marker genes identifying a cluster of rare proliferating T cells (1.3% of total), including two HHV-6 super-expressor cells (82% of HHV-6 UMIs). **(c)** Quantification of all ChIP-seq libraries from CD4+ and CD8+ TIL cultures (PRJNA901909). The abundance of HHV-6 MAPQ 30+ reads is shown with donors stratified by three participating clinical trials. Arrow indicates a high HHV-6 reactivation donor with no matched RNA-seq. **(d)** Single nucleotide variant analysis of Donor 10 and Donor 24 CD8+ ChIP-seq analysis. Shown are allele frequencies of 72 high-confidence single-nucleotide variants that discriminate the viral strains of the two donors.

**Extended Data Fig. 7:**
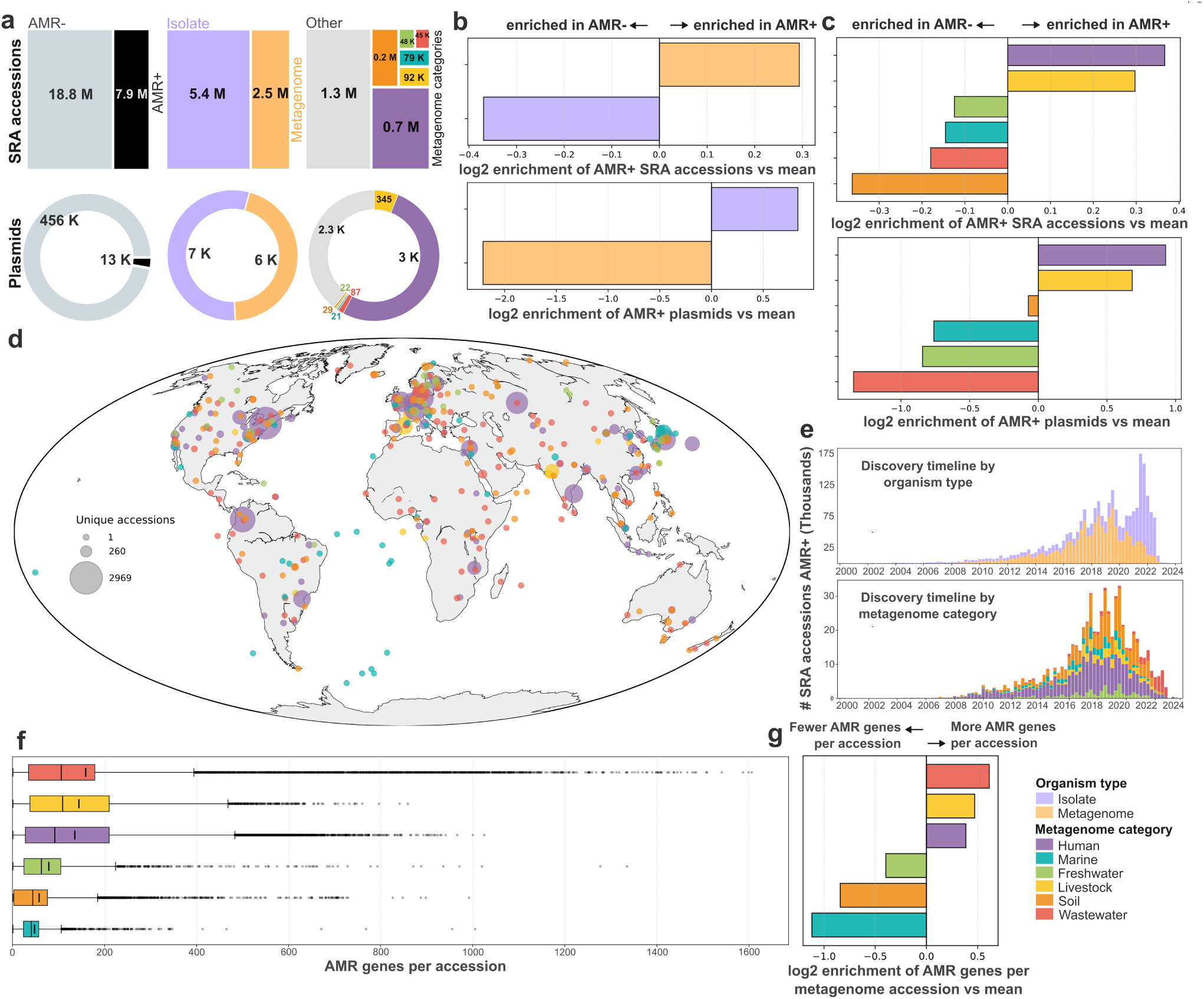
Global distribution of AMR-associated SRA accessions. **(a)** Summary of SRA accessions (top row) and plasmids (bottom row) categorized as AMR-positive (AMR+). First panel, amount of AMR+ vs AMR-samples in the datasets. Second panel, from the AMR+ samples, how many are classified as isolate (purple) or metagenome (yellow) as organism type. Final panel, from the AMR+ metagenome samples, distribution across metagenome categories (human: purple, soil: orange, livestock: yellow, marine: blue, freshwater: green, wastewater: red, other: grey). **(b)** Log2 enrichment of organism type categories in AMR+ datasets versus the average, in SRA accessions (top) and plasmids (bottom), showing relative over- or underrepresentation. Data has been randomly subsampled to avoid bias driven by categories with higher amount of data. **(c)** Log2 enrichment of metagenome categories among AMR+ datasets compared to the mean, for both SRA accessions (top) and plasmids (bottom). Positive values indicate overrepresentation in AMR+ samples. Data has been randomly subsampled to avoid bias driven by categories with higher amount of data. **(d)** Geographic distribution of unique AMR+ SRA accessions across the globe, coloured by metagenome category. Circle size indicates the number of unique accessions per location. **(e)** Temporal trends in AMR gene discovery. Top: Collection date timeline of AMR+ accessions by organism type (isolate: purple, metagenome: yellow). Bottom: Collection date timeline of AMR+ metagenome accessions coloured by metagenome category. **(f)** Distribution of AMR gene counts per accession by metagenome category. **(g)** Log2 enrichment of AMR gene counts per metagenome accession compared to the mean, by metagenome category. Positive values indicate metagenomes with more AMR genes per accession on average. Data has been randomly subsampled to avoid bias driven by categories with higher amount of data. In panels (c), (d), (e) bottom panel, (f), and (g), metagenome category “other” was removed from the analysis.

**Extended Data Fig. 8:**
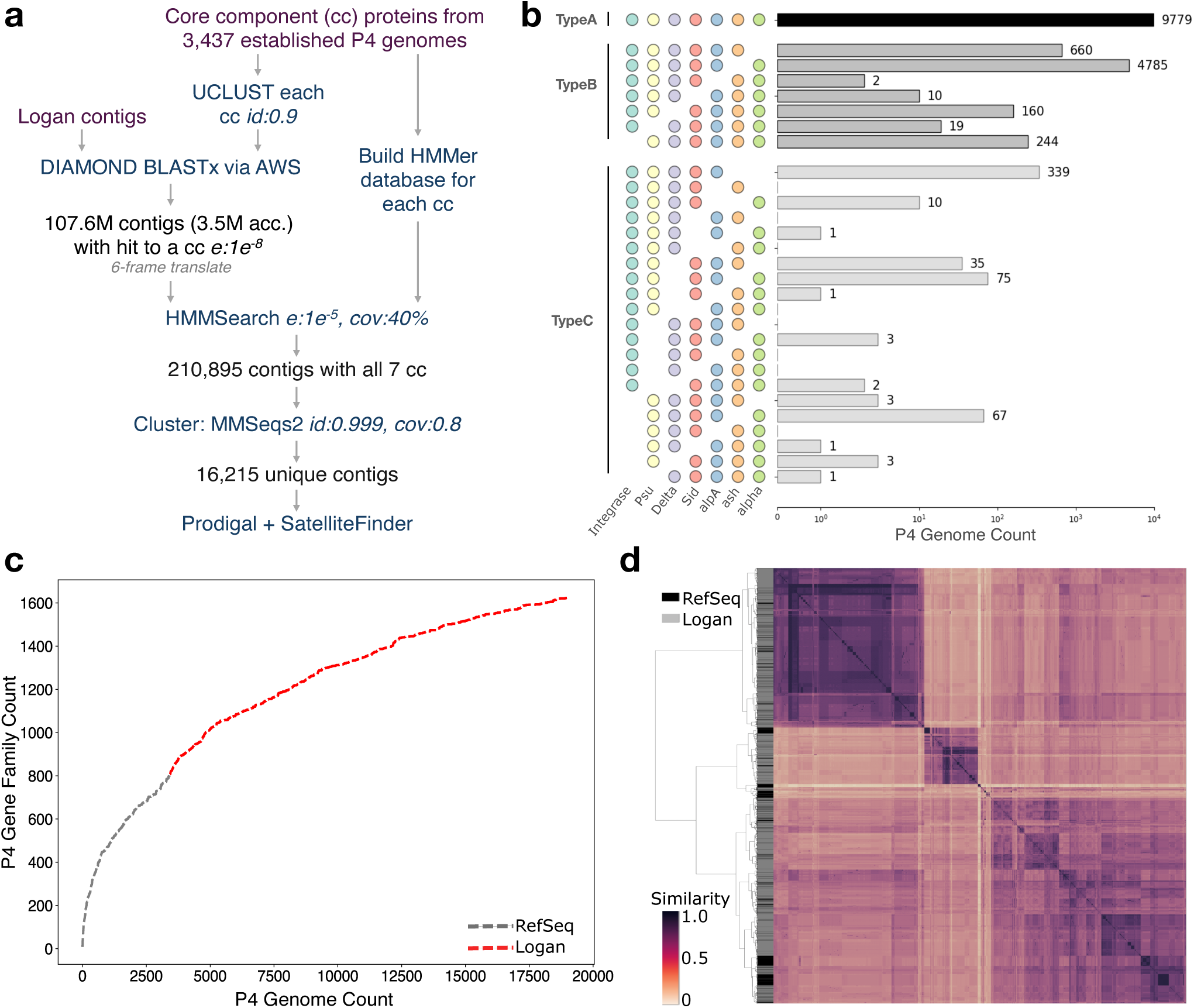
Expansion of P4 phage satellite genetic diversity. **(a)** Pipeline for discovering novel P4 elements. **(b)** Histogram of novel P4 elements binned by SatelliteFinder type. **(c)** Pangenome curve expressing accumulation of gene families clustered at 40% protein identity before (Ref-Seq: Types A, B, and C) and after Logan expansion (Logan: Types A and B). **(d)** Weighted Genome Relatedness Ratio Plot (wGRR) of full proteomes, defined as all proteins found between first and last detected core gene, from before (RefSeq/black) and after (Logan/grey) Logan expansion, where a darker color denotes higher similarity.

